# Predicting Real-life Drinking Scenarios through a Physiological Digital Twin Incorporating Secondary Alcohol Markers

**DOI:** 10.1101/2025.11.14.686953

**Authors:** Henrik Podéus, Christian Simonsson, Gerd Jakobsson, Robert Kronstrand, Elin Nyman, William Lövfors, Gunnar Cedersund

## Abstract

Alcohol consumption poses significant societal challenges, necessitating accurate tools for detecting at-risk drinking. Various biomarkers reflect alcohol intake over different timeframes. *Blood alcohol concentration* (BAC) and *breath alcohol concentration* (BrAC) are commonly used for short-term detection, particularly in forensic contexts such as *driving under the influence of alcohol* (DUIA) cases. Moreover, the rapid kinetics of BAC and BrAC limit their utility in determining the precise timing of intake—an essential factor in legal cases involving defenses like the ‘hipflask’ argument. To address this, secondary metabolites including *ethyl glucuronide* (EtG), *ethyl sulphate* (EtS), and *urine alcohol concentration* (UAC) offer slower, more time-sensitive profiles. Combining these markers could enable a more accurate reconstruction of past alcohol consumption events. Traditionally, mathematical models have been used to help extract information from the dynamics of alcohol-related markers. Existing mathematical models typically focus on primary markers or single secondary markers in isolation. In this study, we present an extended mathematical model that integrates all markers; BAC, EtG, EtS, and UAC into a unified framework, expanding our previous physiological twin model. Our updated model enables personalized simulations of alcohol metabolism and intake timing, which in turn creates the fundament for enhancing forensic assessments and supporting applications where accurate temporal analysis of alcohol consumption is critical. To facilitate accessibility and practical use of this analysis, we have implemented and provided an interactive web application.

## Introduction

There are several problems related to alcohol consumption in society. Therefore, it is important to have accurate screening tools to detect at-risk drinking habits. Today, several markers exist for determining alcohol consumption each with a unique dynamic profile. For long-term alcohol consumption the blood marker *phosphatidylethanol* (PEth) is emerging as a clinical screening tool (1–3). For short-term alcohol consumption, the *blood alcohol concentration* (BAC) or *breath alcohol concentration* (BrAC), are traditionally used (4,5). These short-term markers are often used in forensic analysis of *driving under the influence of alcohol* (DUIA) cases (5,6). While reliable, the fast absorption- and elimination phase of these markers (7,8), and their dependence on anthropometrics (9,10) and paired consumption (11–13), makes it difficult to determine the timing of the alcohol intake. This capability is of importance in judicial cases such as the ‘hipflask’ defence (14–16), where relative precise determination of past events is important. In an effort to more precisely determine the timing of past alcohol intake, new markers of alcohol consumption have been explored (7,17). These markers, *ethyl glucuronide* (EtG), *ethyl sulphate* (EtS), and *urine alcohol concentration* (UAC), have a slower rate of appearance compared to BAC (7). Thus, measuring these metabolites add time-sensitive information of the alcohol metabolism. By overlapping the different profiles of the alcohol metabolism from these markers the ambition is to be able to more accurately determine previous events of alcohol consumption. While promising, to be able to fully extract the information continued in the intersection of these profiles, one needs to employ an analytical tool such as mathematical modelling.

Mathematical modelling has historically been a fundamental tool for estimating the consumption of alcohol, using e.g., Widmark formula (18) or BAC focused models (19–21), typically relaying on the elimination rate of alcohol. With the addition of secondary metabolites, the traditional models become insufficient. For secondary metabolites, mathematical models exist describing the appearance and kinetics of either EtG or EtS following single intakes of alcohol (19,22–24). Furthermore, mathematical models are recently being used as digital twins for describing specific biological functions for single individuals (25–27). We have previously reported such a digital twin model (28), that connects BAC and BrAC profiles with the long-term alcohol blood marker PEth. The model is lacking the connection to EtG, EtS, and UAC. To make use of all information for a e.g., forensic analysis to combat the hipflask’ defence, a common framework simultaneously integrating all primary and secondary markers would be needed. However, no current model exists that integrate all alcohol markers into one cohesive framework.

Herein, we aim to present an updated model extending our previous mathematical model to also describe EtG, EtS, and UAC. In addition, we aim to present how the updated model could be used as a digital twin for describing new data measuring BAC, EtG, EtS and UAC from single individuals.

## Results

We have constructed a mechanistic model (Fig. 1D) describing the dynamics of primary and secondary ethanol metabolites based on our previous work (28). The new model features include: i) addition of the metabolic interactions of EtG and EtS in the liver and the following uptake in the plasma (7,17), ii) introduction of the bladder compartment and the elimination of BAC via the urine, i.e. UAC dynamics (7). Urine based elimination of EtG and EtS was also added. iii) introduction of a tissue compartment and a formulation of *total body water* (TBW) (29), allowing more accurate distribution of BAC, iv) splitting the plasma compartment into a central- and peripheral plasma compartment, to better describe the BrAC dynamics, v) introduction of a diuretic effect through the arginine-vasopressin interactions (30,31), and vi) added an upregulation of enzymatic alcohol elimination when food is present in the stomach (32–34).

**Figure 1:**
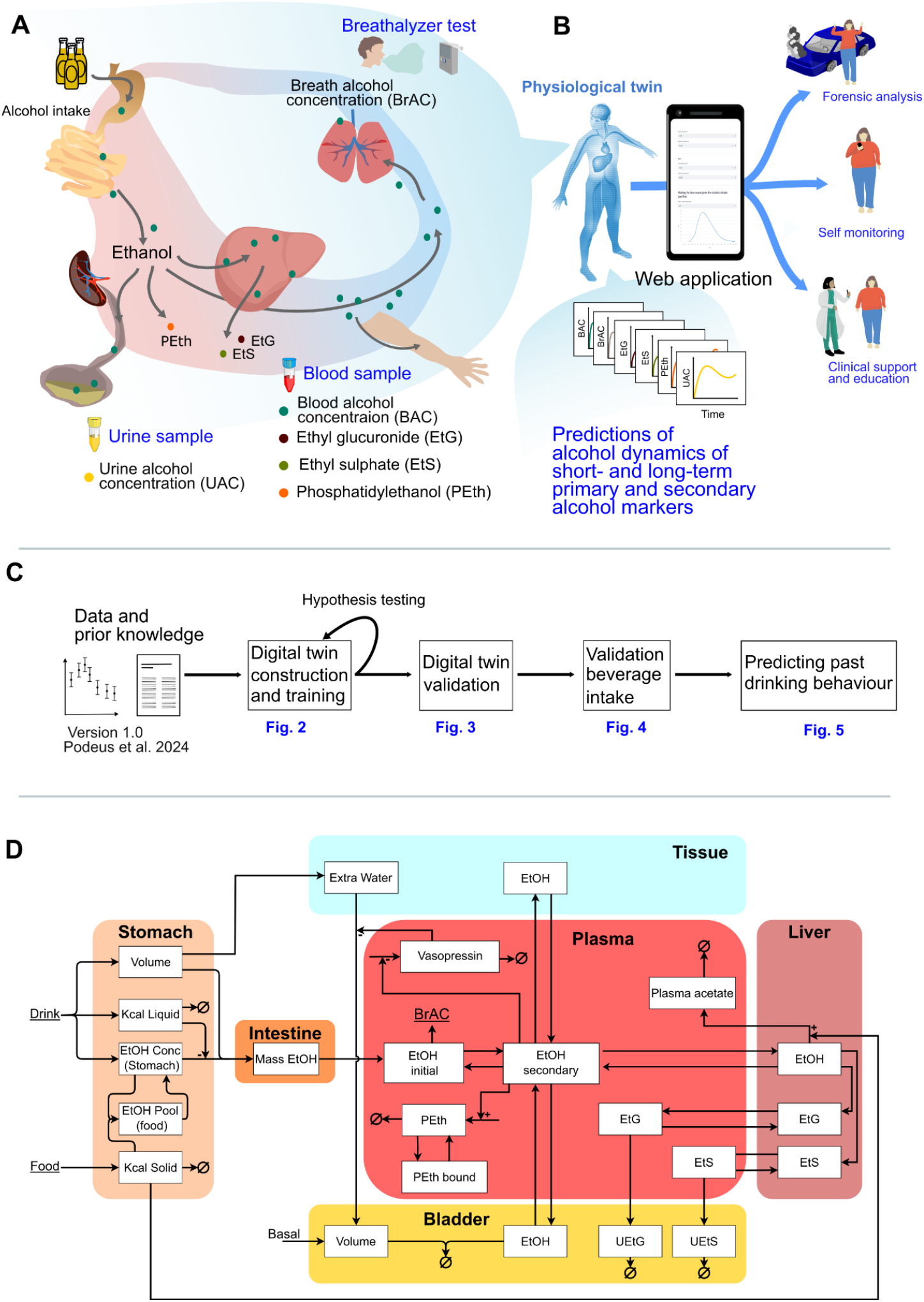
Study overview. **A)** The physiological digital twin presented in this work describes the dynamics of various ethanol markers: *breath alcohol concentration* (BrAC), *blood alcohol concentration* (BAC), *urine alcohol concentration* (UAC), *blood ethyl glucuronide* (EtG), *blood ethyl sulphate* (EtS), and *phosphatidylethanol* (PEth). **B**) The digital twin can offer predictions of the time profiles of the various ethanol markers, which could be valuable in several possible use-cases. For availability the framework is provided in a web application. **C)** In this work we develop a physiological digital twin that incorporates existing knowledge and data of different drinking scenarios from 10 studies. After the model development and validation, the model can make predictions of unknown drinking behaviour from available data samples. **D)** The model consists of: i) a stomach compartment that handles the interactions of alcoholic beverages, non-alcoholic beverages, and food, ii) an intestine compartment from where the alcohol is absorbed into, iii) a plasma compartment from where the ethanol is synthesized into PEth and distributed into either, iv) the bladder where UAC is present, v) the tissue, or vi) the liver where EtG and EtS are synthesized. BrAC is expressed from the plasma ethanol levels in the central compartment.

The model was trained and validated on various published experimental data (12,32,35–41), including dual-drink data from Hoiseth *et al*. (14) and Kronstrand *et al*. (15). The model captures all key behaviours in the estimation data (Fig. 2-3, S1-S3). The model was also validated against independent data, data from Wang *et al*. (41)(Fig. 4). To highlight how the model could be used as a digital twin for all short-term markers, the model was used to predict the alcohol marker profiles (previously unpublished data) of two distinct different individuals (Fig. 5B-I). Lastly, we showcase how the model can be used as a support in, i.e., forensic cases where time profiles of alcohol elimination are investigated – for instance in cases of hip-flask defence (Fig. 5J-Y). Based on this forensic scenario we provide a framework of analysis in an interactive web application (see code availability).

**Figure 2:**
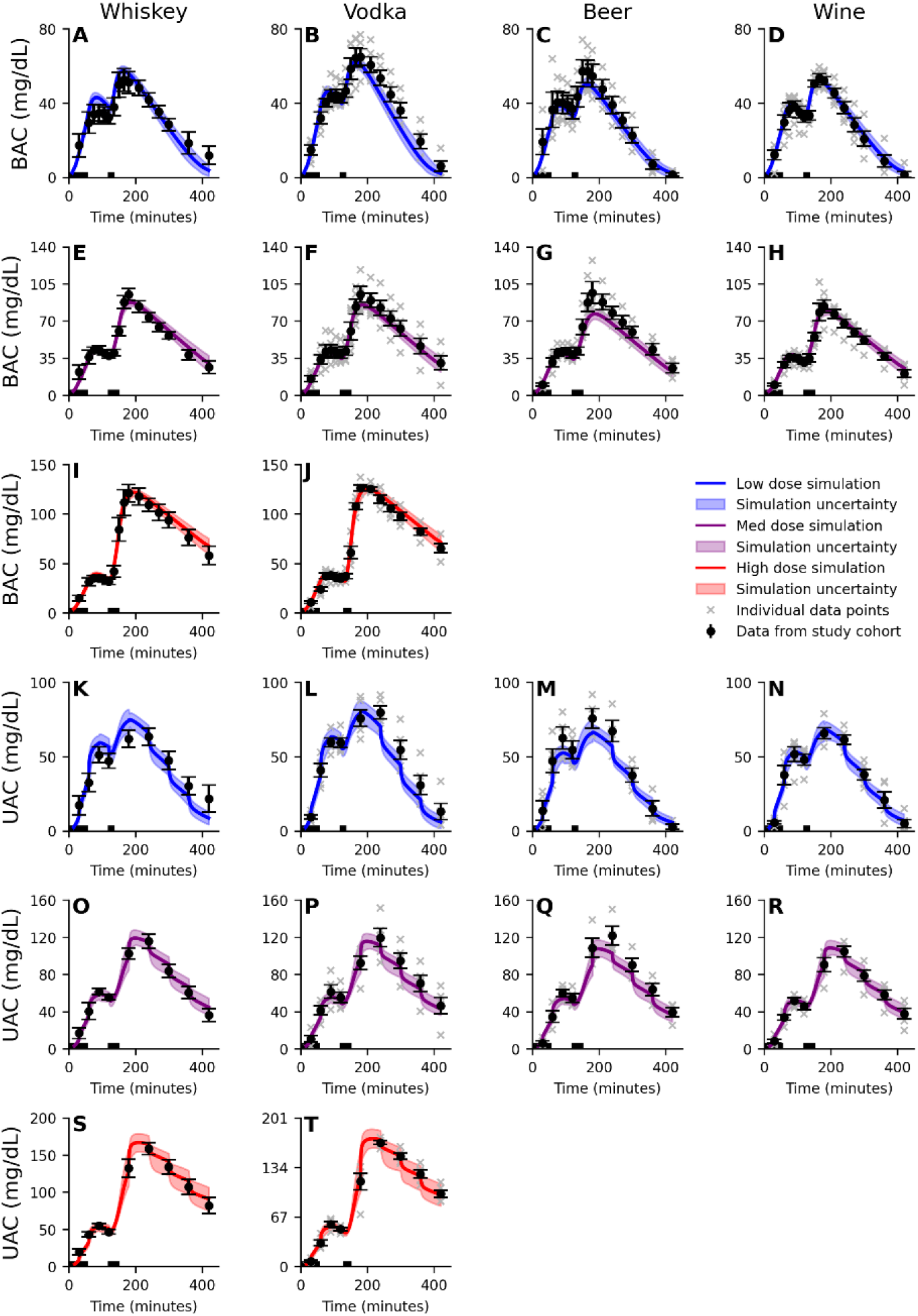
Model agreement to the BAC and UAC data. The solid line is the best model fit (simultaneous agreement to all estimation data), the shaded area is the model uncertainty, the error bars is the *standard error of the mean* (SEM), and the ‘x’ notations indicate the reported individual data points. The model (shaded area) describes the dynamics of the **A-J)** *blood alcohol concentration* (BAC), **K-T)** *urine alcohol concentration* (UAC). All experiments includes a first drink of beer with a 0.51 g/kg dose of ethanol, consumed in 4 periods of 10 minutes over 60 minutes, and a second drink of varying doses, 0.25 g/kg (blue), 0.51 g/kg (purple), or 0.85 g/kg (red), constituted of either; whiskey (A, E, I, K, O, S), vodka (B, F, J, L, P, T), beer (C, G, M, Q), or wine (D, H, N, E). The second drink was consumed over 15 minutes (low dose) or 30 minutes (medium and high dose). The periods where the drinks are consumed are indicated by the black bar over the x-axis. Meals were consumed after the start (300 kcal) of the session and 180 minutes into (500 kcal) the study.

**Figure 3:**
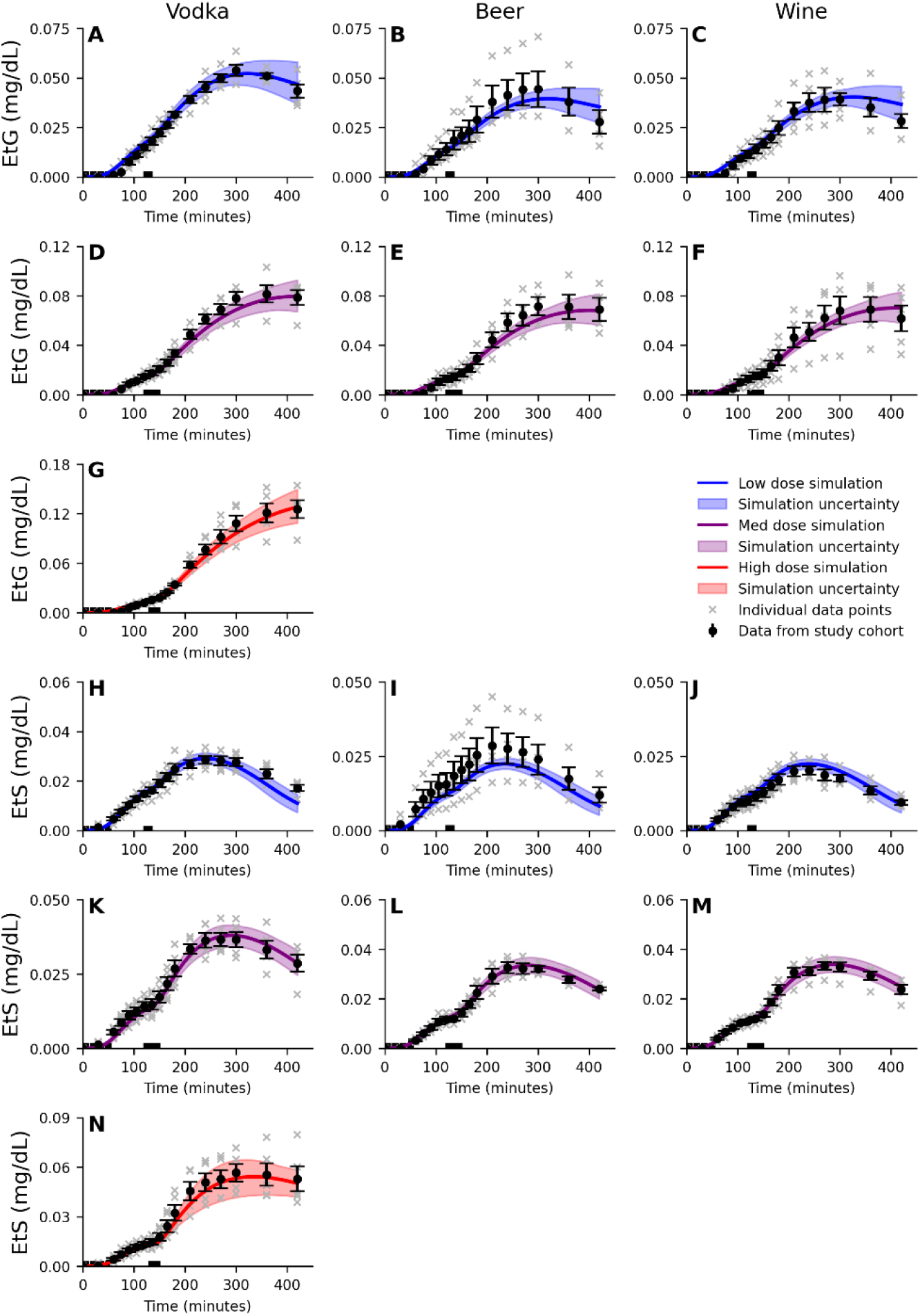
Model agreement to the EtG and EtS data. The solid line is the best model fit (simultaneous agreement to all estimation data), the shaded area is the model uncertainty, the error bars is the *standard error of the mean* (SEM), and the ‘x’ notations indicate the reported individual data points. The model (shaded area) describes the dynamics of the **A-G)** blood *ethyl glucuronide concentration* (EtG), **H-N)** blood *ethyl sulphate concentration* (EtS). All experiments includes a first drink of beer with a 0.51 g/kg dose of ethanol, consumed in 4 periods of 10 minutes over 60 minutes, and a second drink of varying doses, 0.25 g/kg (blue), 0.51 g/kg (purple), or 0.85 g/kg (red), constituted of either; vodka (A, D, G, H, K, N), beer (B, E, I, L), or wine (C, F, J, M). The second drink were consumed over 15 minutes (low dose) or 30 minutes (medium and high dose). The periods where the drinks are consumed are indicated by the black bar over the x-axis. Meals were consumed after the start (300 kcal) of the session and 180 minutes into (500 kcal) the study.

**Figure 4:**
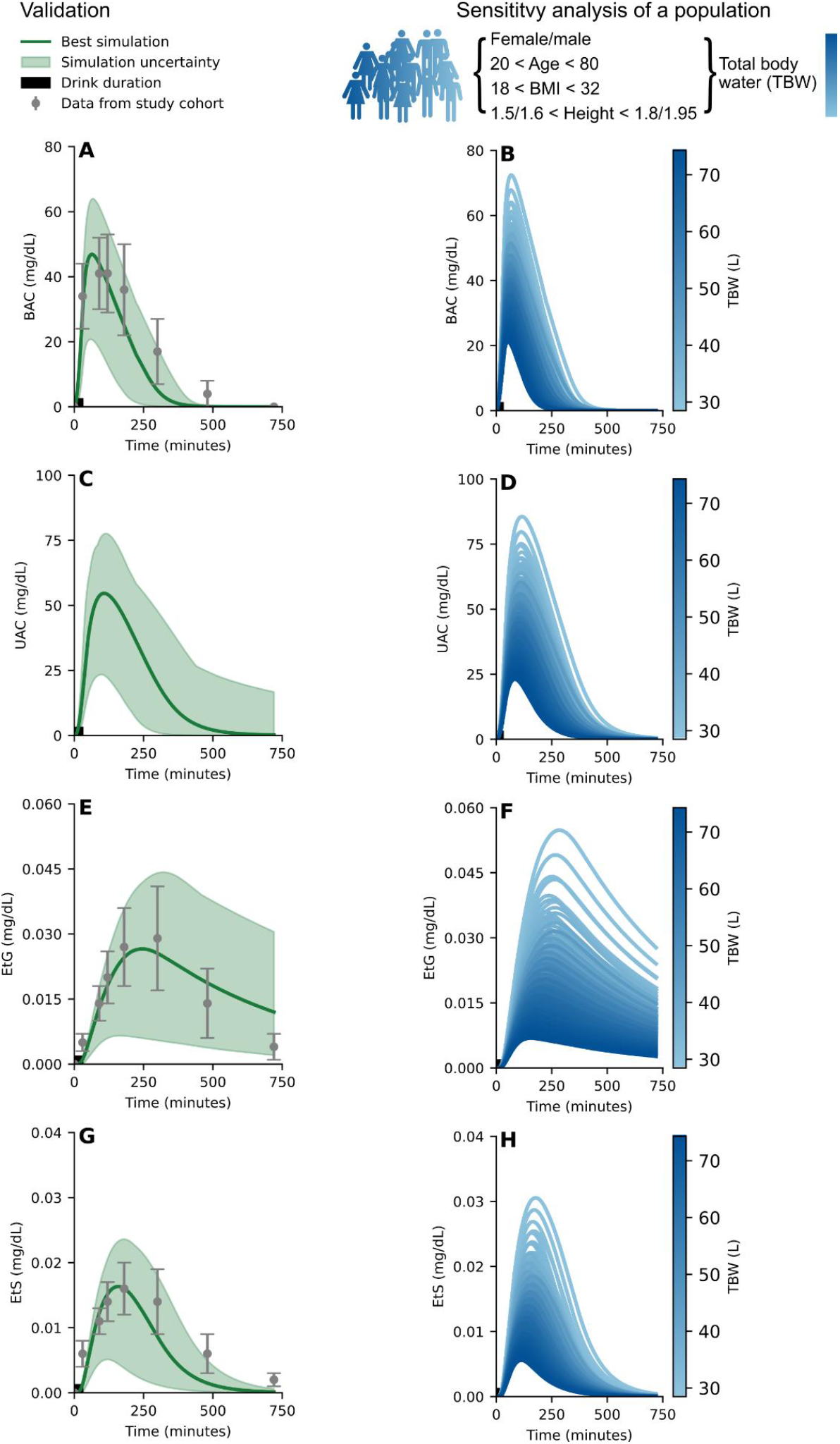
Model validation and sensitivity analysis. The solid line is the simulation using the optimal parameters found for the best model fit to estimation data, the shaded area is the model uncertainty, the grey errorbars indicating the *standard error of the mean* (SEM) for the validation data. The model (shaded area) describes the dynamics of the: **A)** *blood alcohol concentration* (BAC), **C**) *urine alcohol concentration* (UAC), **E)** blood *ethyl glucuronide* (EtG) concentration, and **G)** blood *ethyl sulphate* (EtS) concentration, given a beverage constituted of 0.119 L of 40 v/v% spirits containing 0 kcal, consumed over 30 min (indicated by the black bar on the x-axis and paired with a meal of 500 kcal, consumed over the same 30 min duration. A sensitivity analysis was performed on a population with the anthropometrics ranging between; age 20-80 years, *body mass index* (BMI) 18-32 kg/m^2^, heights of 1.5-1.8 m for females and 1.6-1.95 m for males. This population was challenged with the same drink and food pairing as in **A, C, E**, and **G** and the model simulations of the population are presented as the blue gradient for **B)** BAC, **D)** UAC, **F)** EtG, and **H)** EtS. The blue gradient corresponds to the volume of *total body water* (TBW) for the subjects.

**Figure 5:**
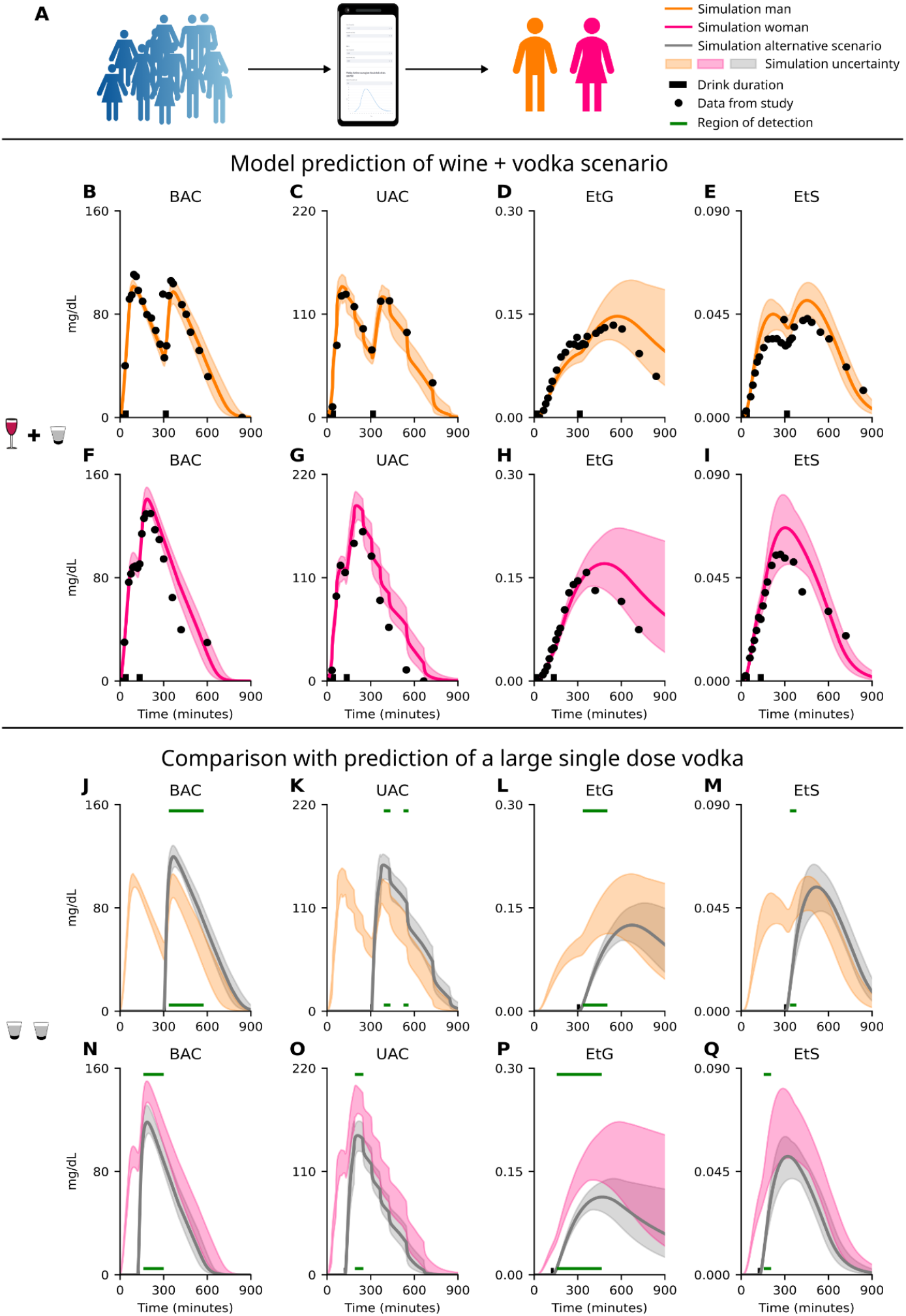
Model predictions of individuals in forensic scenarios. **A)** Illustration showing that the model is trained on a mean population behaviour and is now used to predict individual behaviours, a male (orange) and a female (pink). **B-I)** Model prediction of the four markers; **B, F)** *blood alcohol concentration* (BAC), **C, G)** *urine alcohol concentration* (UAC), **D, H)** blood *ethyl glucuronide* (EtG) concentration, and **E, I)** blood *ethyl sulphate* (EtS) concentration. The drinking challenge for the male and female were consumption of an initial drink containing 0.85 g/kg ethanol constituted of 13 v/v% wine over four 10 minutes block for a total of 60 minutes and a second drink containing 0.51 g/kg ethanol constituted of 20 v/v% vodka consumed over 30 minutes. The solid line represents the best model solution from the model training and the area is the model uncertainty. The data points are represented by the black circles, and the black bar indicates the period of alcohol consumption. **J-Q)** An *in-silico* drinking scenario for; **J-M)** the male and **N-Q)** the female, were evaluated. The alternative drinking challenge was a single dose of 1.0 g/kg ethanol constituted of 40 v/v% vodka for both the male and female. The model prediction is shown for the four different markers: **J, N)** BAC, **K, O)** UAC, **L, P)** EtG, and **M, Q)** EtS. The grey solid line represents the best model solution from the model training and the grey area is the model uncertainty. The model prediction for the wine and vodka drinking challenge (**B-I**) for the male is shown as reference for the male (orange **J-M**) and the female (pink, **N-Q**). The horizontal green lines indicate the regions where the model predicts differences between the two drinking challenges for the respective individual (male and female).

### The model can describe EtG, EtS and UAC dynamics

The new developed model was simultaneously trained on data from 10 studies (12,14,15,32,35–40). As already shown in our previous study (28), there is a good agreement between the model and all included studies of single-drink intake (12,32,35–40), see Fig. S1-S3. To also include data on a sequential drink scenario, and EtG, EtS and UAC, we extended the model training with data presented by Kronstrand *et al*. (15) and Hoiseth *et al*. (14). This new data presents the behaviour of BAC (Fig. 2A-J), UAC (Fig. 2K-T), EtG (Fig. 3A-G), and EtS (Fig. 3H-N) in response to two sequential drinks (‘x’, single data points and means with error bars). All different drinking configurations in these two studies, included a first drink of beer with the alcohol content of 0.51 g/kg body weight, followed by a second drink of either: whiskey (Fig. 2A, 2E, 2I, 2K, 2O, 2S), vodka (Fig. 2B, 2F, 2J, 2L, 2P, 2T, 3A, 3D, 3G, 3H, 3K, 3N), beer (Fig. 2C, 2G, 2M, 2Q, 3B, 3E, 3I, 3L), or wine (Fig. 2D, 2H, 2N, 2R, 3C, 3F, 3J, 3M). The colour represents the dose of the second drink (blue - low dose of 0.25 g/kg, purple - medium dose of 0.51 g/kg, and red - high dose of 0.85 g/kg). The best model simulation is presented with solid lines and the model uncertainty as a shaded area. There was a good simultaneous agreement with all training data, confirmed by a visual assessment and a χ^2^ test. The χ^2^ test statistic was 642.74 which was lower than the cutoff (T_χ2_=735.51, for n=674).

### Model validation of alcohol markers

Following the model training, the model was validated against an independent data set from Wang *et al*. (41), see Fig. 4A, 4E, 4G. The subjects in the study consumed 0.119 L of vodka (40 v/v%), together with a meal of 500 kcal, over 30 minutes and BAC, EtG, and EtS were measured in blood at several time points over 720 minutes. This independent study was first simulated with the model and then compared with the data from Wang *et al*. (41). The model validation also passed the χ^2^ test and the model uncertainty is shown as the shaded area (Fig. 4A, 4C, 4E, 4G). The χ^2^ test statistic was 28.04 which was lower than the cutoff (T_χ2_=31.41, for n=20). The model was further evaluated by performing a sensitivity analysis where the anthropometrics were varied to represent the variability of a population (age 20-80 years, *body mass index* (BMI) 18-32 kg/m^2^, and a height of 1.5-1.8 m for females and 1.6-1.95 m for males). The model simulations of this population, given the same drinking scheme presented by Wang *et al*. (41), are presented as the blue-green gradient (Fig. 4B, 4D, 4F, 4H), where the gradient indicate the TBW in litre (L) of the subjects, which is influenced by all the anthropometric variables (see Eq. 9). These simulations highlight the robustness of the model, as the model behaviour is qualitatively preserved between the individuals over the whole population, which is of importance in applications of model personalization. We next investigated how well the validated model functions in additional use cases, including personalized predictions for single individuals.

### Model prediction of individual alcohol marker profiles

Following the validation and sensitivity analysis, the model’s ability to make personalized prediction was evaluated using newly collected data from single individuals (Fig. 5A). The data includes BAC, UAC, EtG, and EtS for sequential drinks and the raw data is available in the Supplementary Information, see “S1 Raw prediction data”. The study was approved by the Swedish Ethical Review Authority, 2023-02640-01, see methods. The data describes two individuals, a male (Fig. 5B-E, orange), body weight 106.6 kg, age 28 years, height 1.86 m, TBW 66.2 L, blood volume 6.4 L, and female (Fig. 5F-I, pink), bodyweight 62.7 kg, age 22 years, height 1.69 m, TBW 37.4 L, blood volume 4.0 L. The model prediction, shaded area, aligns well with the experimental data, black circles, (Fig. 5B-I) indicating that although the model is trained on a mean population behaviour it can be used for predictions of individuals. The drinking challenge for the male and female were consumption of an initial drink containing 0.85 g/kg ethanol constituted of 13 v/v% wine over four 10 minutes block for a total of 60 minutes and a second drink containing 0.51 g/kg ethanol constituted of 20 v/v% vodka consumed over 30 minutes. Meals (300 kcal, 500 kcal, and 500kcal) were consumed at 90-, 180-, and 540 minutes for the male, and 30-, 180-, and 540 minutes for the female.

As a showcase of the model’s ability to accurately predict distinct drinking challenges, and as such also separate between similar challenges, we present an alternative scenario. This *in-silico* scenario (Fig. 5J-Q) replaces both drinks with a single vodka challenge (1.0 g/kg ethanol) for the male (Fig. 5J-M) and female (Fig. 5N-Q). This scenario represents plausible alternative drinking challenge. For the *in-silico* scenario, the BAC profile closely aligns with segments of the experimental data (Fig. 5J and 5N) and would be hard to differentiate from the actual drinking challenge (Fig. 5B and 5F). This highlights a current limitation in, for instance, forensic analysis of alcohol elimination. By making use of the model’s ability to simultaneously describe all markers, we can distinguish between wide range of plausible drinking challenges and offer insights of deviating elimination profiles, indicated by the green lines (Fig. 5J-Q). This example highlights how our physiological digital twin could be used as a decision support in forensic analysis.

## Discussion

Herein, we present a physiological twin that can describe the dynamics of different markers of alcohol consumption: BAC (Fig. 2, S2), UAC (Fig. 2), EtG (Fig. 3), and EtS (Fig. 3), and PEth (Fig. S3). The model was validated using independent data (Fig. 4A, E, G), and the robustness was evaluated using a sensitivity analysis (Fig. 4B, D, F, H). The validated model was able to make personalized predictions of new individual data generated from dual drinking challenge (Fig. 5B-I). Furthermore, we show how the model predictions can differentiate between similar drinking behaviours by simulating an *in-silico* drinking challenge (Fig. 5J-Q). By evaluating the model predictions, the model can be used to differentiate between different claims of alcohol consumption. This capability could be utilized as a decision support tool in forensic analysis, where on need to determine the time profile of alcohol elimination.

### Model estimation, validation, and robustness analysis

The model was trained and validated to a total of 10 different study datasets, which include different markers of ethanol consumption, including: BAC, UAC, EtG, EtS, and PEth (Fig. 2,3, S1-S3). An overview of the data is given in the Supplementary Information, “4 Usage of experimental data”. The model could explain all estimation data (Fig. 2-3, S1-S3) and validation data (Fig. 4) to a satisfactory level. Altogether, the model sufficiently describes the dynamics of these ethanol consumption markers following consumption of beverages (single and dual) of different volumes, concentrations, time of consumption, in combination with food, and for individuals with different anthropometric data.

While the model passed a χ2-test for all estimation data, it is worth pointing out some aspects of the data that the model did not fully capture. Firstly, the model had some difficulty to describe the peak BAC following the consumption of the second drink of the medium dose of vodka (Fig. 2F) and beer (Fig. 2G), data presented by Hoiseth *et al*. (14). In contrast, the model describes the medium dose well for whiskey (Fig. 2E), data presented by Kronstrand *et al*. (14), and wine (Fig. 2H), even though the dose of ethanol is the same as for beer and vodka. This difference points towards an inconsistency in the model explanation of the rate of gastric emptying. This discrepancy could stem from how the drink was consumed during the study, and what the drink was paired with, an aspect that we have explored in our previous work (28). In the Hoiseth *et*.*al*. study (14) and the Kronstrand *et*.*al*. study (15), the consumption of non-alcoholic beverages was not recorded, and as such no alcohol-free drinks were included in the model inputs. Alcohol-free drinks were freely available during the experiment and were therefore likely consumed and could thus be the reason for the difference we observe in the medium dose. Secondly, we observe the same differences in peak UAC between vodka (Fig. 2P) and beer (Fig. 2Q), a too low peak, versus whiskey (Fig. 2O) and wine (Fig. 2R), a well described peak. This is expected as urine concentration is highly correlated with the blood concentration. Finally, one can note that the EtS concentration in the case of the low dose of beer (Fig. 3I) are underestimated. In this case, the experimental data is higher compared to the medium dose of beer (Fig. 3L) which is not mirrored in the BAC experimental data (Fig. 2C, G). The model does not have the ability to explain this difference in EtS synthesis, given similar BAC concentrations, and will therefore underestimate EtS for a low dose of beer. This behaviour could possibly be caused by a single subject with higher EtS synthesis (see grey ‘x’ markers, in Fig. 3I) which influences the mean behaviour not seen in the other groups.

There are also some qualitative differences between model predictions and corresponding validation data. For the validation, the model predicts a faster elimination of BAC than seen in the experimental data (Fig. 4A). As the model describes the rate of appearance of BAC well, one is inclined to believe that the drink is estimated well. Although the drinking challenge is probably not fully representative to the one consumed in the Wang *et al*. study (41), as the subject were tasked to consume a 40 v/v% beverage within 30 minutes and did likely not consume it evenly over this 30 minutes period (which we assume due to no other information available). Secondly, because of the rapid elimination of BAC we also observe a too fast clearance of EtS (Fig. 4G). Thirdly, we observe that the EtG instead is eliminated too slowly (Fig. 4E). This is likely a result due to the lack of EtG data showing the full elimination time-period in our model estimation data set. Since the model is not trained on the full elimination profile of EtG – the model predicts a slow elimination profile in the later, unknown, elimination phase. Finally, we can see that the model behaviour is preserved when challenging individuals within a population (age 20-80 years, body mass index (BMI) 18-32 kg/m^2^, and a height of 1.5-1.8 m for females and 1.6-1.95 m for males) with the same drinking challenge (0.119 L 40 v/v% over 30 minutes) (Fig. 4B, D, F, H). As expected, the drink has a higher influence on smaller individual (less TBW, light blue) and a lower influence on larger individuals (more TBW, dark blue).

### Data considerations

Due to the lack of information in the different included studies some assumptions have been made and should be discussed. Firstly, some of the studies did not report anthropometrical data of the subjects or details regarding several aspects of the experimental setup e.g., reported drinking time. Also, for some datasets mean values were calculated in a mixed cohort including both female and male participants (14,15,38,39). To take this into consideration we opted to estimate the mean blood volume based on the sex distribution. Also, for some studies the caloric content was not reported, leading to some assumptions regarding the consumed beverage. The caloric content is of importance as the gastric emptying module is the major contributor to the ethanol rate of appearance in plasma. For details regarding these assumptions see Supplementary Information “3 Input estimation”. Secondly, the EtG and EtS concentrations were reported as zero until they exceeded the detection limit in the Hoiseth *et al*. study (14). Since the subjects in the different groups passes the detection limit at different time points, we choose to exclude the data from the time points where all subjects had not reached the detection limit. In practice, EtS was part of the estimation data when time > 30 minutes and EtG was part of the estimation data when time > 75 minutes. Thirdly, as non-alcoholic beverages were available during the experiments in the Hoiseth *et al*. and Kronstrand *et al*. studies (14,15), we assumed that the participants consumed an equal volume of non-alcoholic beverages as the volume of 40 v/v% spirits (due to the large volume of hard spirits). This effectively doubles the volume and halves the concentration of the drink. Fourth, the study protocol in the Hoiseth *et al*. and Kronstrand *et al*. studies (14,15) included a blood sample every 15 minutes and a urine sample every 30 minutes during the drinking challenge. We consider the effects of this as the first drink being consumed in four periods of ten minutes, leaving five minutes for data collection. Lastly, we decreased the drinking window for the high vodka challenge in Hoiseth *et al*. (14). This is due to the BAC experimental data lagging and barely increasing 15 minutes into the reported drinking window. Compared to the high whiskey challenge in Kronstrand *et al*. (15), where BAC increases substantially faster, we assume that this discrepancy is due to hesitation of drinking. As such, we implement the vodka drinking challenge to start ten minutes later, reduced the drinking window to twenty minutes.

### The predictive capability of drinking behaviour of the model

The validated physiological twin was able to predict the profiles of the different alcohol markers, BAC (Fig. 5B, 5F), UAC (Fig. 5C, 5G), EtG (Fig. 5D, 5H), and EtS (Fig. 5E, 5I), for two new individuals – a large male (Fig. 5B-E) and a small female (Fig. 5F-I). While the predictions are generally impressive and points towards the robustness of the model, especially considering that only the anthropometric parameters in the model were tuned for these predictions, it is worthwhile to mention some deviations. While the BAC and UAC predictions are closely in line with the data (black circles, Fig. 5), EtG and EtS are a bit more deviating. This is not too surprising since the model was not able to fully describe the variability in the EtG and EtS levels during model training. This indicates that the model does not incorporate all the information needed to describe the inter-variability of EtG and EtS between individuals (42–44), resulting in the offset of the simulation and generally a larger uncertainty (shaded area). By investigating the variability in enzymatic expression of UDP-glucuronosyltransferase (43,45) and sulfotransferase (46), one could better identify the individual differences and reduce the model variability. While this would be useful information and improve the physiological detail of the model, it is not feasible to always accompany the blood sample with an enzymatic test. Instead, the tuning would need to rely on further model development through larger data sets.

### The potential of physiological twins as support in forensic cases

The model was evaluated with an *in-silico* scenario (Fig. 5J-Q), constructed to be similar to the real drinking challenge (Fig. 5B-I). This scenario was constituted of a single larger drink of vodka (Fig. 5J-Q) and produced a model behaviour that described segments of the experimental data. But importantly, there are deviating behaviour between this scenario and the model prediction of the ‘real’ drinking challenge (Fig. 5B-I). The regions that showcase the deviating behaviour is indicated by the green line (Fig. 5J-Q) and highlight where the model predictions of the original drinking challenge (orange for male and pink for female) and the *in-silico* scenario (grey for both male and female) deviates from each other. Only regions after the consumption of the last drink had ended are highlighted. As can be seen, the different markers (BAC, UAC, EtG, and EtS) have different time intervals where the predicted behaviour deviates between the model predictions. This is of importance as it allows us to intersect the time profiles of the different markers to evaluate if the given scenario could have generated the experimental data. The different profiles of the markers also allow for a more robust evaluation of the claimed drinking challenge, allowing the twin to identify small differences in the scenarios.

This capability could be of use in i.e. forensic analysis of DUIA cases, where one needs to determine if the claimed drinking challenge is in accordance with the gathered data samples. Our model framework is well suited to offer support in such scenarios and to aid in the determination of the plausibility of a claimed scenario. To fully utilize the model framework, one would first need to investigate the sensitivity of the analysis depending on the delay between data sampling and the end of drinking and how the reliability of the different alcohol markers is affected with different number of data samples. Overall, the personalization capabilities of our modelling framework could improve the accuracy of the forensic analysis and aid in distinguishing between similar drinking challenges.

## Conclusion

To summarize, we have presented a physiological twin that accurately describe a wide variety of drinking scenarios, including multiple markers of alcohol consumption: BAC, UAC, EtG, and EtS. We showcase that our physiological twin can make accurate predictions of unknown individuals and that the robust model predictions can be consulted to distinguish between similar drinking scenarios. Our model framework could with further validation offer support in forensic analysis of DUIA cases where current methodologies are insufficient to combat e.g., the hip-flask defence.

## Methods

Within this section the model equations are detailed, see Equations 2-9. The full model structure is shown in figure 1D.

Before detailing the equations, an example of an *ordinary differential equation* (ODE) is described. A typical ODE used in this work looks similar to Eq. (1).

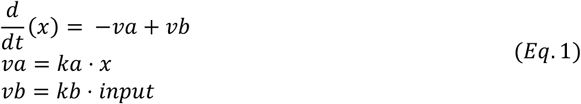

Here, *x* is a state in the model, *va* and *vb* are reaction rates, *ka* and *kb* are rate-determining parameters, and *input* is some input to the state. In other words, the amount of the state *x* is decreased by the reaction *va* with the speed *ka* and increased by the reaction *vb* with the speed *kb* depending on some input *input*.

### Model description

This model was built upon our previous work, see Podéus *et al*. (28). In the following sections, the changes and additions to the model are reported. The full model structure is available in the supplementary code and the iterative model development is described in detail in the Supplementary Information, see “5 Changelog of rejected model structures”.

#### Gastric emptying

The gastric emptying module was updated to the following format.

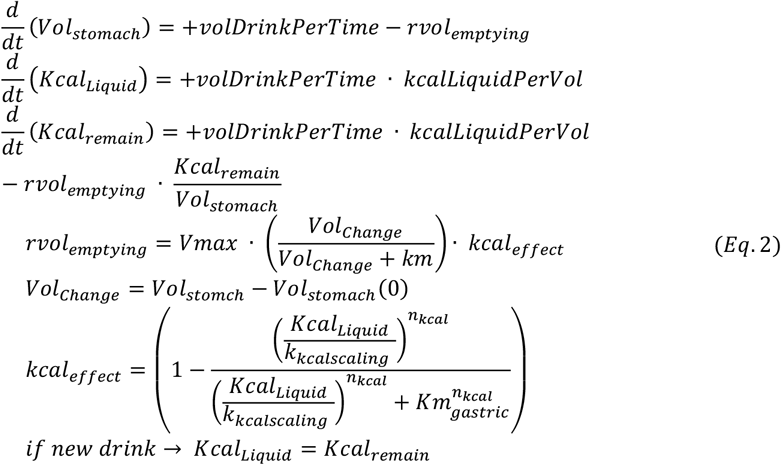

Here, the gastric volume is governed by the consumption of new liquids, *volDrinkPerTime*, and passing of liquid to the intestines, *rvol*_*emptying*_. The caloric contents of the stomach are governed by the new *Kcal*_*remain*_ state that: i) keeps track of the incoming kcal, *kcalLiquidPerVol*, ii) the emptying rate of kcal, and iii) updates the *Kcal*_*Liquid*_ state once a new drink is consumed.

#### Blood compartment split into central and peripheral

To account for the difference in rate of appearance in BrAC and BAC, the blood compartment was split into a central compartment, where BrAC is measured, and a peripheral compartment, where BAC is measured. The ethanol is absorbed into the central compartment, from the intestines, via *rEtOH*_*uptake*_ as a mass (mg) and diluted into the blood volume of the central compartment. The ethanol then diffuses between the central and peripheral compartments, via *r*_*Circulation*_. The downstream reactions, diffusion into the tissue *r*_*Tissue Peripheral*_, uptake to the liver *r*_*Liver*_, and transportation to, *r*_*Urine*_, and from, *r*_*Urine return*_, the bladder are feed from the peripheral compartment.

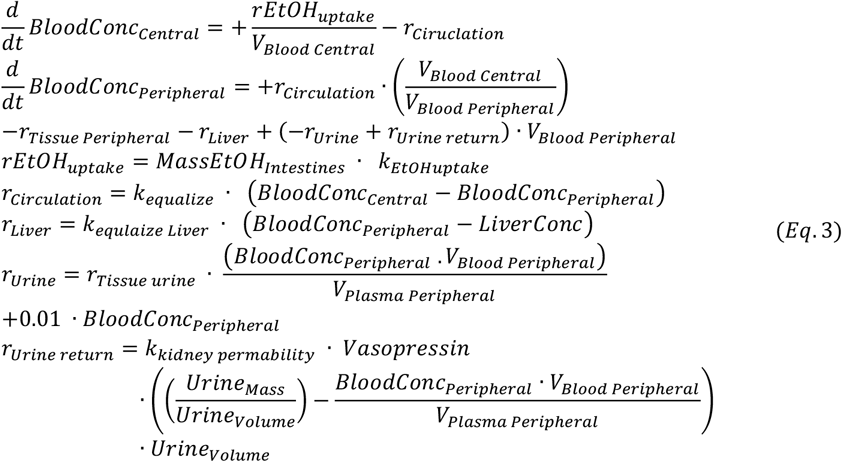

Where the blood volumes are based on Nadler’s equation (47), and the plasma volume is estimated using a parameter, *k*_*blood plasma ratio*_, representing the water contents of the blood.

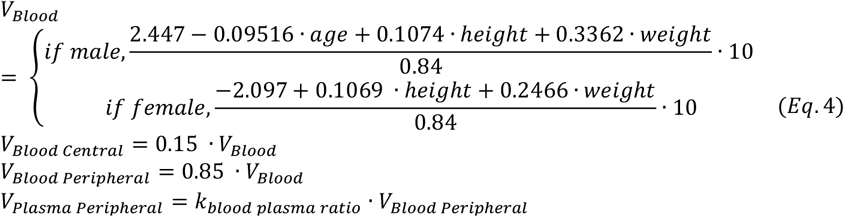

#### Dynamics of EtG and EtS

EtG and EtS are synthesised from the breakdown of BAC in the liver compartment, diffused into the peripheral blood compartment, and eliminated through the urine.

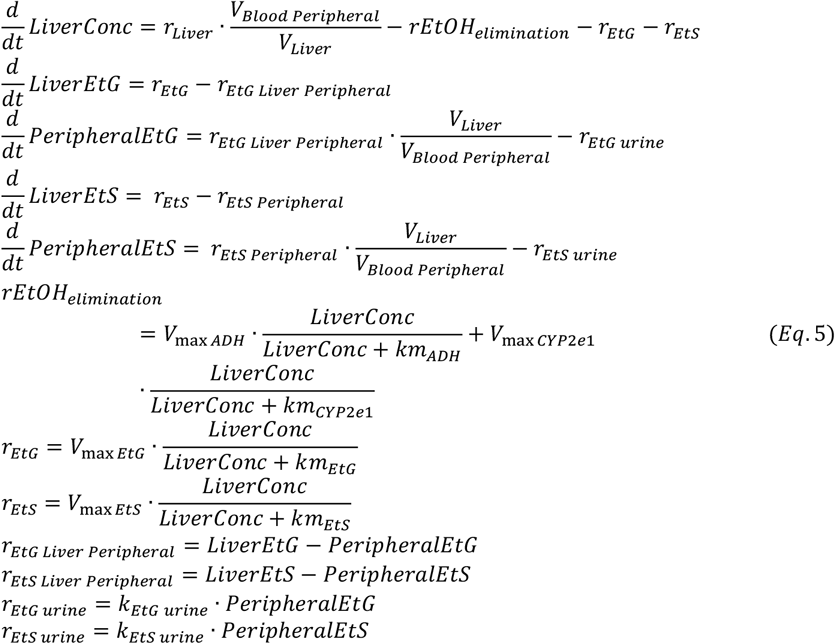

Where the volume of the liver is described according to Vauthey’s formula (48).

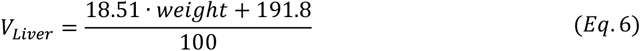

#### Dynamics of UAC

The urine is handled as a mass and volume, instead of a concentration, as it makes it easier to keep track of consumed liquid that continuously enter the bladder.

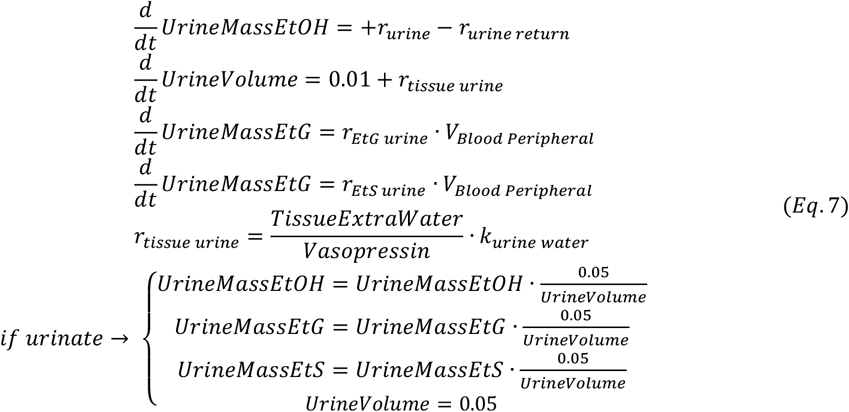

Where 0.05 dL represent the residual volume in the bladder after urination (49).

#### Dynamics of tissue and vasopressin

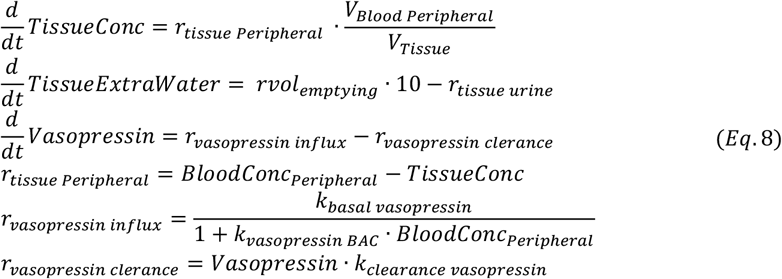

Where the tissue volume is described as the difference between the TBW, Watson *et al*. (29), and the other volumes divided into separate compartments in the model.

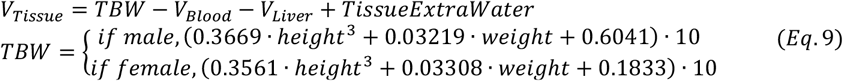

### Initial values of the model

It was assumed, in the model, that the person has no residual alcohol in the system. Furthermore, it was assumed that the model starts in a fasted state, with no kcal in the system, that the residual volume in the stomach was 0.001 L, and that the residual volume in the bladder was 0.05 dL. The initial value of *Vasopressin* was estimated by assuming mass balance in the first time point. In the case of the Javors experiments (38), the basal values of *PEth* and *PEthBound* were also estimated from the estimated parameter values assuming mass balance in the first time point. Otherwise, they were assumed to be 0. The initial values used are given in Table 1.

**Table 1:** Initial values of the model.

| State | Initial values |
| --- | --- |
| $Vol_{Stomach}$ | $1.0 \cdot 10^{-3}$ |
| $Kcal_{Liquid}$ | 0.0 |
| $Kcal_{remain}$ | 0.0 |
| $MaxKcal_{Solid}$ | 0.0 |
| $Kcal_{Solid}$ | 0.0 |
| $EtOH_{Pool}$ | 0.0 |
| $ConcEtOH_{Stomach}$ | 0.0 |
| $MassEtOH_{Intestines}$ | 0.0 |
| $BloodConc_{Central}$ | 0.0 |
| $BloodConc_{Peripheral}$ | 0.0 |
| $TissueConc$ | 0.0 |
| $LiverConc$ | 0.0 |
| $LiverEtG$ | 0.0 |
| $PeripheralEtG$ | 0.0 |
| $LiverEtS$ | 0.0 |
| $PeripheralEtS$ | 0.0 |
| $Plasma_{Acetate}$ | 0.0 |
| $TissueExtraWater$ | 0.0 |
| $Vasopressin$ | *estimated* |
| $UrineMassEtOH$ | 0.0 |
| $UrineVolume$ | 0.05 |
| $UrineMassEtG$ | 0.0 |
| $UrineMassEtS$ | 0.0 |
| $PEth$ | *estimated* |
| $PEth_{Bound}$ | *estimated* |
| $timeElapsed$ | 0.0 |

### Model parameter values

This section gives the optimal parameter values for the connected model when estimated to the estimation dataset (columns *θ*_*est*_^*^). Furthermore, the bounds used in the optimization for all parameters are also given (columns lower bound and upper bound), see Table 2. *km*_*ADH*,_ *km*_*CYP*2*E*1_, and *km*_*Gastric*_ were given bounds reported in literature (50).

**Table 2:** Parameter values and parameter estimation bounds.

| Parameter | $\theta_{est}^*$ | lower bound | upper bound |
| --- | --- | --- | --- |
| VmaxGastric | 0.10028847533601028 | $1 \cdot 10^{-5}$ | $1 \cdot 10^5$ |
| KmGastric | 0.8115471295053232 | $2.766 \cdot 10^1$ | $1.844 \cdot 10^3$ |
| k_kcalscaling | 0.026009077044725923 | $1 \cdot 10^{-5}$ | $1 \cdot 10^5$ |
| km_kcal | 7219.025648168511 | $1 \cdot 10^{-5}$ | $1 \cdot 10^5$ |
| n_kcal | 2.9350431505014307 | $1 \cdot 10^{-1}$ | $4 \cdot 10^0$ |
| k_poolIn | 0.049215056963481905 | $1 \cdot 10^{-5}$ | $1 \cdot 10^5$ |
| k_poolOut | 0.00011026963373147828 | $1 \cdot 10^{-7}$ | $1 \cdot 10^5$ |
| VmaxADHSto | 383.0316652068957 | $1 \cdot 10^{-5}$ | $1 \cdot 10^5$ |
| KmADHSto | 1750.8565753331234 | $2.766 \cdot 10^2$ | $1.844 \cdot 10^3$ |
| k_EtOHuptake | 0.05085931578651339 | $1 \cdot 10^{-5}$ | $1 \cdot 10^5$ |
| k_equalize | 3.364285096106452 | $1 \cdot 10^{-5}$ | $1 \cdot 10^5$ |
| k_equalize_liver | 0.5664934932515018 | $1 \cdot 10^{-5}$ | $1 \cdot 10^5$ |
| VmaxADH1 | 7.490115599723829 | $1 \cdot 10^{-5}$ | $1 \cdot 10^5$ |
| VmaxCYP2E1 | 0.7715882956131286 | $1 \cdot 10^{-5}$ | $1 \cdot 10^5$ |
| KmADH1 | 2.13180346563429 | $9.22 \cdot 10^{-1}$ | $9.22 \cdot 10^0$ |
| KmCYP2E1 | 45.98540429542478 | $3.688 \cdot 10^1$ | $4.61 \cdot 10^1$ |
| k_food_clearance | 0.1350242774593947 | $1 \cdot 10^{-5}$ | $1 \cdot 10^5$ |
| k_acetate | 1.1650870599405063 | $1 \cdot 10^{-5}$ | $1 \cdot 10^5$ |
| k_blood_plasma_ratio | 0.6871729866586396 | $1 \cdot 10^{-1}$ | $1 \cdot 10^0$ |
| k_urine_water | 5.961711956546037 | $1 \cdot 10^{-5}$ | $1 \cdot 10^5$ |
| k_basal_vasopressin | 0.3849587409310818 | $1 \cdot 10^{-5}$ | $1 \cdot 10^5$ |
| k_vasopressin_BAC | 74740.43401768291 | $1 \cdot 10^{-5}$ | $1 \cdot 10^5$ |
| k_clearance_vasopressin | 0.0006626630440471179 | $1 \cdot 10^{-5}$ | $1 \cdot 10^5$ |
| k_kidney_permability | 2.5215624280668018e-05 | $1 \cdot 10^{-5}$ | $1 \cdot 10^5$ |
| VmaxEtG | 0.6461457040480787 | $1 \cdot 10^{-5}$ | $1 \cdot 10^5$ |
| KmEtG | 29854.322456892394 | $1 \cdot 10^{-5}$ | $1 \cdot 10^5$ |
| VmaxEtS | 0.011876595138349037 | $1 \cdot 10^{-5}$ | $1 \cdot 10^5$ |
| KmEtS | 414.40740948670106 | $1 \cdot 10^{-5}$ | $1 \cdot 10^5$ |
| k_EtG_urine | 0.0024879684263205454 | $1 \cdot 10^{-5}$ | $1 \cdot 10^5$ |
| k_EtS_urine | 0.014152433882270748 | $1 \cdot 10^{-5}$ | $1 \cdot 10^5$ |
| k_PEth | 0.13959486810293995 | $1 \cdot 10^{-7}$ | $1 \cdot 10^3$ |
| k_PEth_clearance | 0.001460712263112933 | $1 \cdot 10^{-7}$ | $1 \cdot 10^3$ |
| k_PEth_bind | 0.07805739912357477 | $1 \cdot 10^{-7}$ | $1 \cdot 10^3$ |
| k_PEth_release | 0.002851044320061577 | $1 \cdot 10^{-7}$ | $1 \cdot 10^3$ |

### Model inputs

This section lists the input values the model needs; see Table 3. A detailed overview of all the inputs provided to the model for each dataset is provided in the Supplementary Information, see “3 Input estimations”.

**Table 3:**
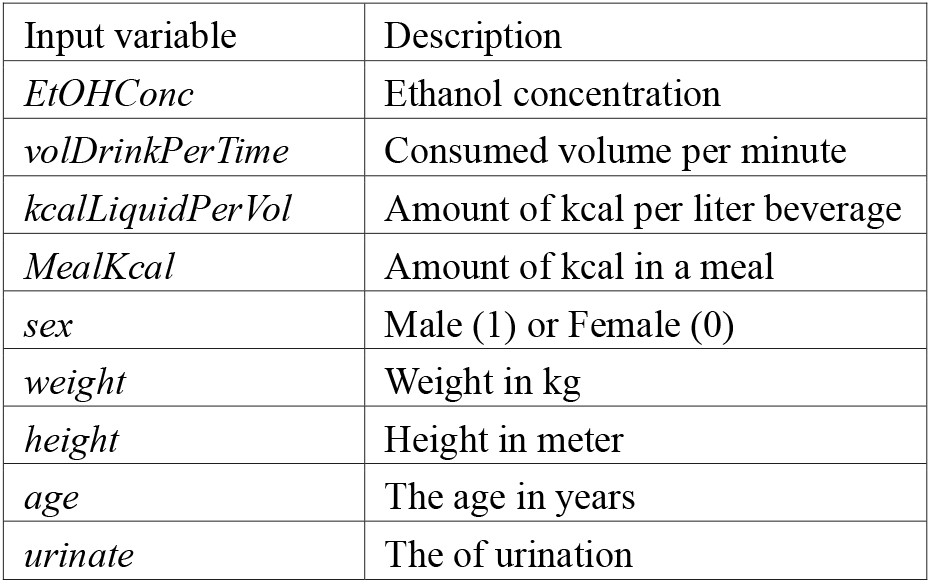
Input information to the model.

### Model outputs

This section lists the model outputs, and the scaling performed. *yBrAC*_*g201L*_ rescales the plasma concentration of ethanol *BloodConc*_*Central*_ into breath concentration using a linear correlation observed between BrAC and BAC measurements by Skaggs et al. (51). The additional division of 1000 is to go from *g* to *mg*. In Javors *et al*. (38), the blood concentration was estimated from the breathalyzer test and as such the *yBrAC*_*gdL*_ uses *BloodConc*_*Central*_ and scales it with 1000 to get g/dL. *yAcetate* is divided with 10.2 convert the concentration unit *mg/dL* to *mM. yUAC* is calculated from the mass of EtOH in the urine and the volume of urine in the bladder.

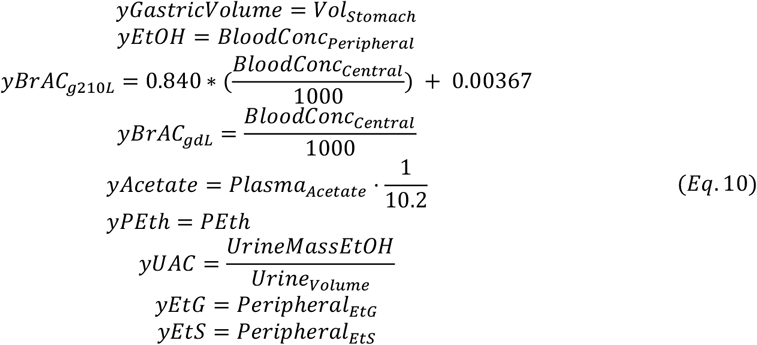

### Parameter estimation

All model analysis was performed using Python 3.10.4 and plotting using Python 3.12.3 (52). The simulations was carried out using the SUND toolbox (53). For model parameter estimation the dual annealing (54) and differential evolution (55,56) algorithms, provided by SciPy (57), were used.

Parameter estimation was done by quantifying the model performance, using the model output ŷ to calculate the traditional weighted least squares cost function defined as

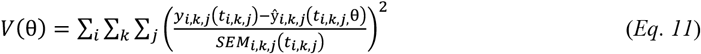

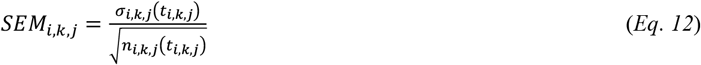

where, θ is the model parameters; *y*_*i,k,j*_(*t*_*i,k,j*_) is the measured data from a study *i*, and from on type of measure *k*, at time point *j*; ŷ_*i,k,j*_(*t*_*i,k,j*_, θ) is the simulation value for a given experiment setup *i*, type of measure *k*, and time point *j*. SEM is the standard error of the mean, which is the sample standard deviation, *σ*_*i,k,j*_(*t*_*i,k,j*_) divided with the square root of the number of repeats, *n*_*i,k,j*_(*t*_*i,k,j*_) at each time point. The value of the cost function, *V*(θ), is then minimized by tuning the values of the parameters, typically referred to as parameter estimation.

To evaluate the new model, a *χ*^2^-test for the size of the residuals, with the null hypothesis that the experimental data have been generated by the model, and that the experimental noise is additive and normally distributed was performed. In practice, the cost function value was compared to a *χ*^2^ test statistic,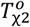. The test statistic *χ*^2^ cumulative density function,

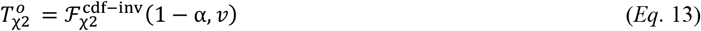

where 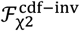 is the inverse density function; and α is the significance level (α = 0.05, was used) and *v* is the degrees of freedom, which was equal to the number of data points in the estimation dataset (674 in total, all timepoints over all experiments). In practice, the model is rejected if the model cost is larger than the *χ*^2^-threshold 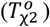.

### Uncertainty estimation

The model simulation uncertainty was gathered as proposed in (58) and is visulised as the uncertainty areas in the figures. The model uncertainty is estimated by dividing the problem into multiple optimization problems, with one problem per model property 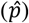. In this work, the property 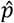 corresponds to either a simulation at a specific time point *j, ŷ*(*t*_*j*,_θ), or a parameter value 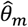. Each problem is solved by maximizing and minimizing the property value, while satisfying that the cost (*V*(θ)) is below the *χ*^2^-threshold 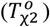. By identifying themaximal and minimal value of the model property (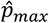 and 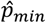), the boundary values of the property uncertainty area are found. Mathematically, this operation for the parameter values is formulated as,

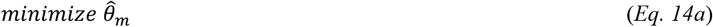

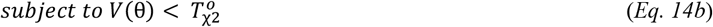

where 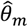 is minimized to find the lower value of the parameter, while also satisfying that the cost (*V*(θ)) is below the *χ*^2^-threshold 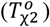. To find the upper bound of the uncertainty area the problem is maximized instead. In practice, the constraint (Eq 4b) can be relaxed into the objective function as a L1 penalty term with an offset if 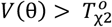.

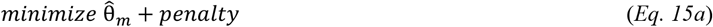

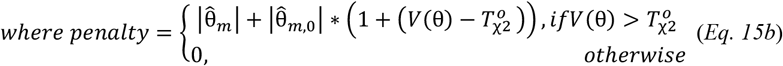

Here, the penalty is scaled with the initial value of the parameter, 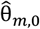 and the offset between the cost and the *χ*^2^-threshold 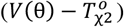. To maximize the parameter 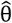, and thus finding the upper bound of the uncertainty area, the problem is solved as a minimization problem. This is done by substituting 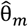 with 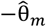 in the objective function. To solve the problem for the model simulation at a specific time point, *ŷ*(*t*_*j*_, θ), the problem is formulated as follows,

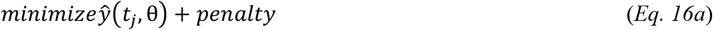

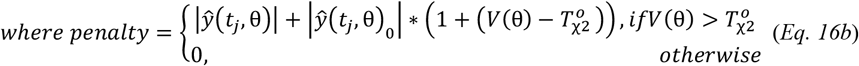

Here, the penalty is scaled with the initial value of the parameter, *ŷ*(*t*_*j*_, θ)_0_, and the offset between the cost and the *χ*^2^-threshold 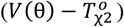. To maximize the model simulation at time point *j*, (*ŷ*(*t*_*j*_, θ), and thus finding the upper bound of the uncertainty area, the problem is solved as a minimization problem. This is done by substituting *ŷ*(*t*_*j*_, θ) with −*ŷ*(*t*_*j*_, *θ*) in the objective function.

### The experimental data used for the modelling

This work incorporates a wide variety of data for the model estimation and validation - details of these data is given below.

The gastric emptying module was evaluated with three studies from Okabe *et al*.; the first explored the effect of caloric content (35), the second the influence of caloric density (36), and the third the effect of alcoholic calories (12).

A variety of studies observing the BAC levels were included. The effect of a meal was studied in Jones *et al*. (32). Mitchell *et al*. investigated the effect of different alcoholic compositions (37). Sarkola *et al*. studied the BAC and acetate response following one drink (39). Frezza *et al*. compared the effect between females and males (40). Javors *et al*. investigated the BrAC and PEth levels after a drink (38). Kronstrand *et al*. (15) and Hoiseth *et al*. (14) investigate the BAC response to double dosing of different alcohol types and volumes.

The alcohol marker UAC was collected by Kronstrand *et al*. (15) and Hoiseth *et al*. (14). EtG and EtS markers were reported by Hoiseth *et al*. (14).

Wang *et al*. presented the data used for the validation analysis, which was constituted of BAC, EtG, and EtS measurements (41).

We collected new data from two individuals during a sequential drink intervention study performed by the National Board of Forensic Medicine.. Written consent was obtained from all participants, and the study was approved by the Swedish Ethical Review Authority (2023-02640-01).

## Abbreviations

AC: blood alcohol concentration
MI: body mass index
rAC: breath alcohol concentration
DUIA: driving under the influence of alcohol
EtG: ethyl glucuronide
EtS: ethyl sulfate
SEM: standard error of the mean
PEth: phosphatidylethanol
TBW: total body water
UAC: urine alcohol concentration

## Data availability statement

All data used for model estimation and validation can be accessed from the original publications. We provide all model related data files and parameter values in our public code repository, https://github.com/Podde1/alcohol-secondary-metabolites.git, with a permanent copy available at Zenodo (DOI: https://doi.org/10.5281/zenodo.17609970).

## Code availability statement

All related scripts and data files are provided in our GitHub repository (https://github.com/Podde1/alcohol-secondary-metabolites.git) with a permanent copy available at Zenodo (DOI: https://doi.org/10.5281/zenodo.17609970). Additionally, a user interface for the model implemented as a web application is provided. This application is available at https://alcohol.streamlit.app/, with the source code available from our GitHub repository (https://github.com/willov/alcohol_app) - with a permanent copy available at Zenodo (https://doi.org/10.5281/zenodo.17609892).

## Acknowledgements

The computations were enabled by resources provided by the National Supercomputer Centre (NSC), funded by Linköping University.

The authors acknowledge financial support from: GC acknowledges support from the Swedish Research Council (2023-03186, 2023-05460), VINNOVA (VisualSweden), the Horizon Europe project STRATIF-AI (101080875), and ALF (RÖ-1001928). GC and WL acknowledge support from the Exploring Inflammation in Health and Disease (X-HiDE) Consortium - a strategic research profile at Örebro University funded by the Knowledge Foundation (20200017). EN acknowledges support from the Swedish Research council (2019-03767), and the Swedish Fund for Research without Animal Experiments (S2021-0008, F2022-02). WL acknowledges support from the Area of Strength e-Health at Linköping University and Region Östergötland. WL and GE acknowledge support from the Strategic Research Area in Forensic Sciences. GJ and RK acknowledges support from the strategic research area in Forensic Science at Linköping University (SoFo 2022-09) funded by the National Board of Forensic Medicine and Linköping University.

The funders had no role in study design, data collection and analysis, decision to publish, or preparation of the manuscript.

## Author Contributions

**Conceptualization:** HP, GJ, CS, EN, RK, and WL. **Formal model analysis:** HP, CS, and WL. **Data curation:** HP, GJ, CS, RK, and WL. **Visualization:** HP, CS, and WL. **Supervision:** GC, EN, RK, GJ, and WL. **Funding acquisition:** GJ, GC, EN, and WL.

All authors were involved in writing and reviewing the manuscript. All authors read and approved the final manuscript.

## Competing Interests

The authors declare no competing interests.

## Figure legends

**Figure S1:**
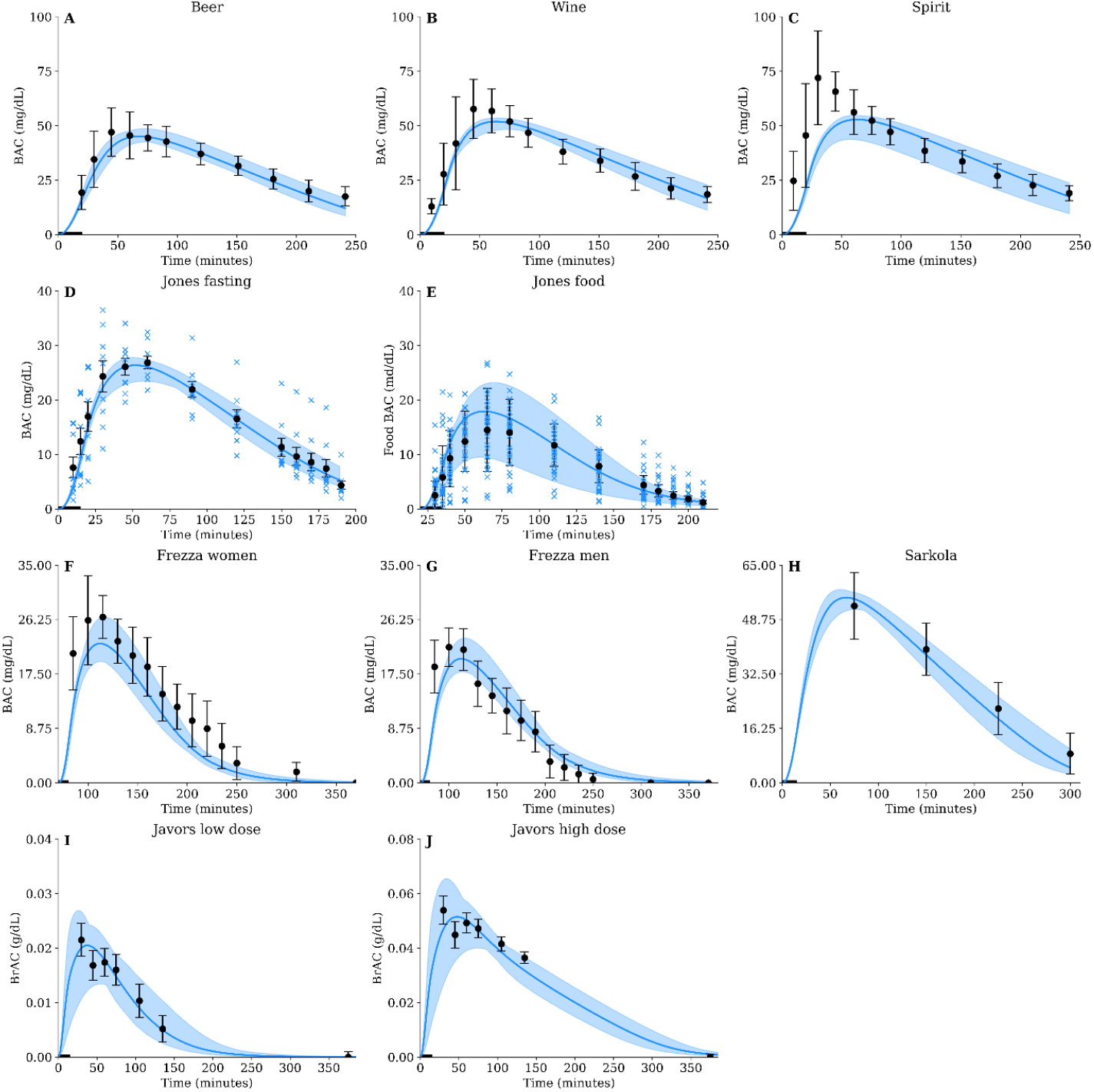
Model agreement to ethanol data for various interventions. The solid line is the best model fit, the shaded area is the model uncertainty, the black boxes are the mean experimental data value with the error bars indicating the *standard error of the mean* (SEM), and individual data points are marked with a “x”. The alcohol was consumed orally through different beverages in various alterations **A)** 1L of 5.1 v/v % beer (133 kcal) was consumed over 20 min, **B)** 0.42L of 12.5 v/v % wine (56 kcal) was consumed over 20 min, **C)** 0.26L of 20 v/v % spirit blend (43 kcal) was consumed over 20 min, **D)** 0.14L of 20.0 v/v % spirit blend (51 kcal) over 15 min, **E)** 0.14L of 20.0 v/v % spirit blend (51 kcal) over 15 min after a meal constituted of 700 kcal, **F)** plasma ethanol levels in women after consumption of 0.19L of 12 v/v % spirit blend (28 kcal) over 10 min after eating a meal constituted of 555 kcal, **G)** plasma ethanol levels in men after consumption of 0.22L of 12 v/v % spirit blend (33 kcal) over 10 min after eating a meal constituted of 555 kcal, **H)** plasma ethanol levels from consumption of 0.48L of 10.0 v/v % spirit blend (149 kcal) over 15 min. Estimated *blood alcohol concentration* (BAC) levels from *breath alcohol concentration* (BrAC) estimates were reported from **I)** consumption of 0.72L of 3.25 v/v % spirit blend (251 kcal) over 15 min, **J)** 0.71L of 6.5 v/v % spirit blend (247 kcal) was consumed over 15 min. In **A-D** and **H-J** the beverage was consumed in a fasting state, and in **E**-**G** in combination with food. In **A-H** the absolute change in BAC was measured and in **I-J** the absolute change in estimated BAC levels derived from BrAC measurements.

**Figure S2:**
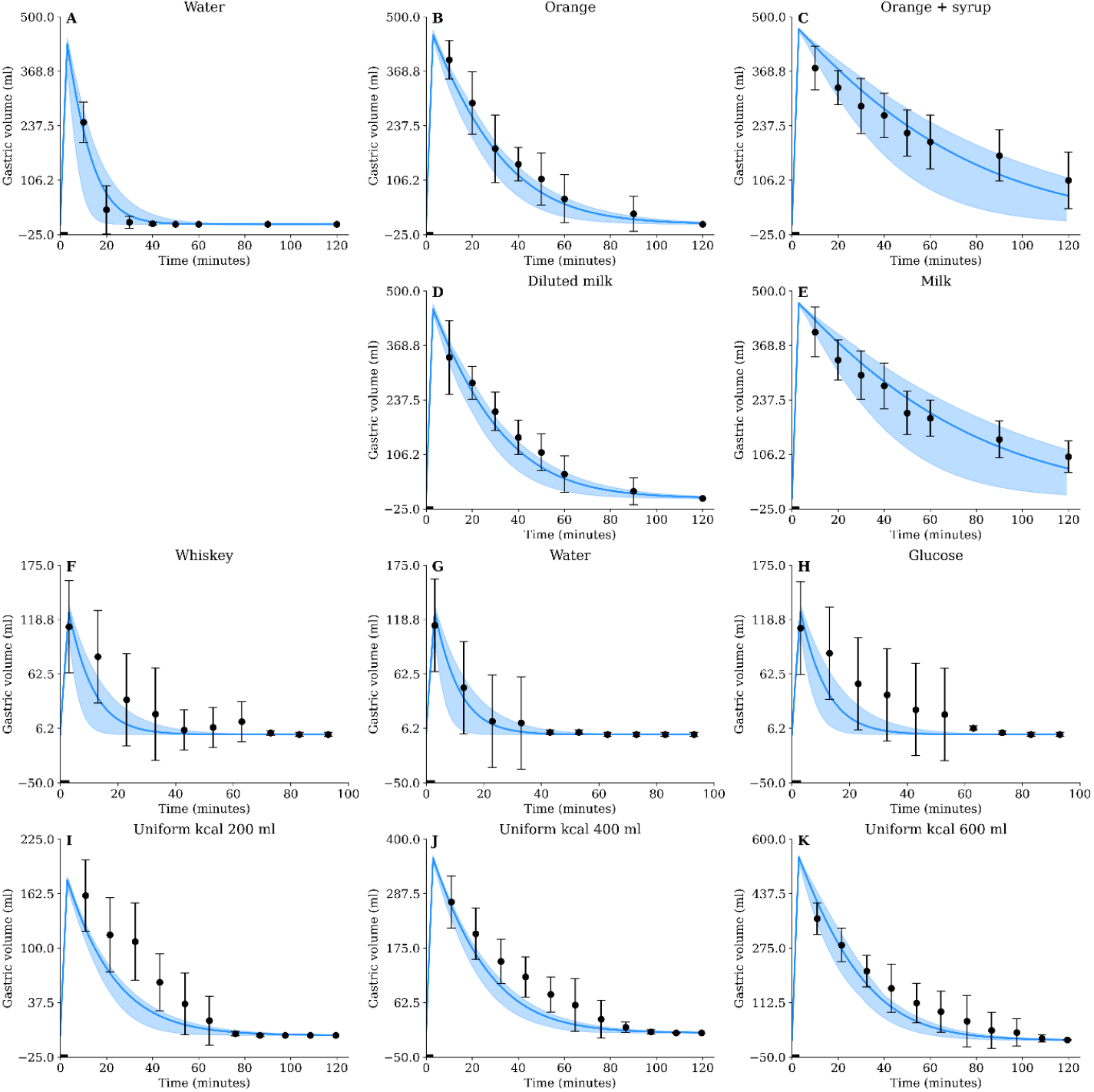
Model agreement to gastric emptying data. The solid line is the best model fit, the shaded area is the model uncertainty, the black boxes are the mean experimental data value with the error bars indicating the *standard error of the mean* (SEM). The model (solid lines) describes the clearance of different beverages from the stomach (blue markers with error bars). In all experiments, the beverage was consumed over 3 minutes and was constituted of the following: **A)** 500 mL containing 0 kcal, **B)** 500 mL containing 220 kcal, **C)** 500 mL containing 329 kcal, **D)** 500 mL containing 220 kcal, **E)** 500 mL containing 329 kcal, **F)** 150 mL containing 0 non-ethanol kcal, **G)** 150 mL containing 0 kcal, **H)** 150 mL containing 67 kcal, **I)** 200 mL containing 200 kcal, **J)** 200 mL containing 400 kcal, and **K)** 600 mL containing 200 kcal.

**Figure S3:**
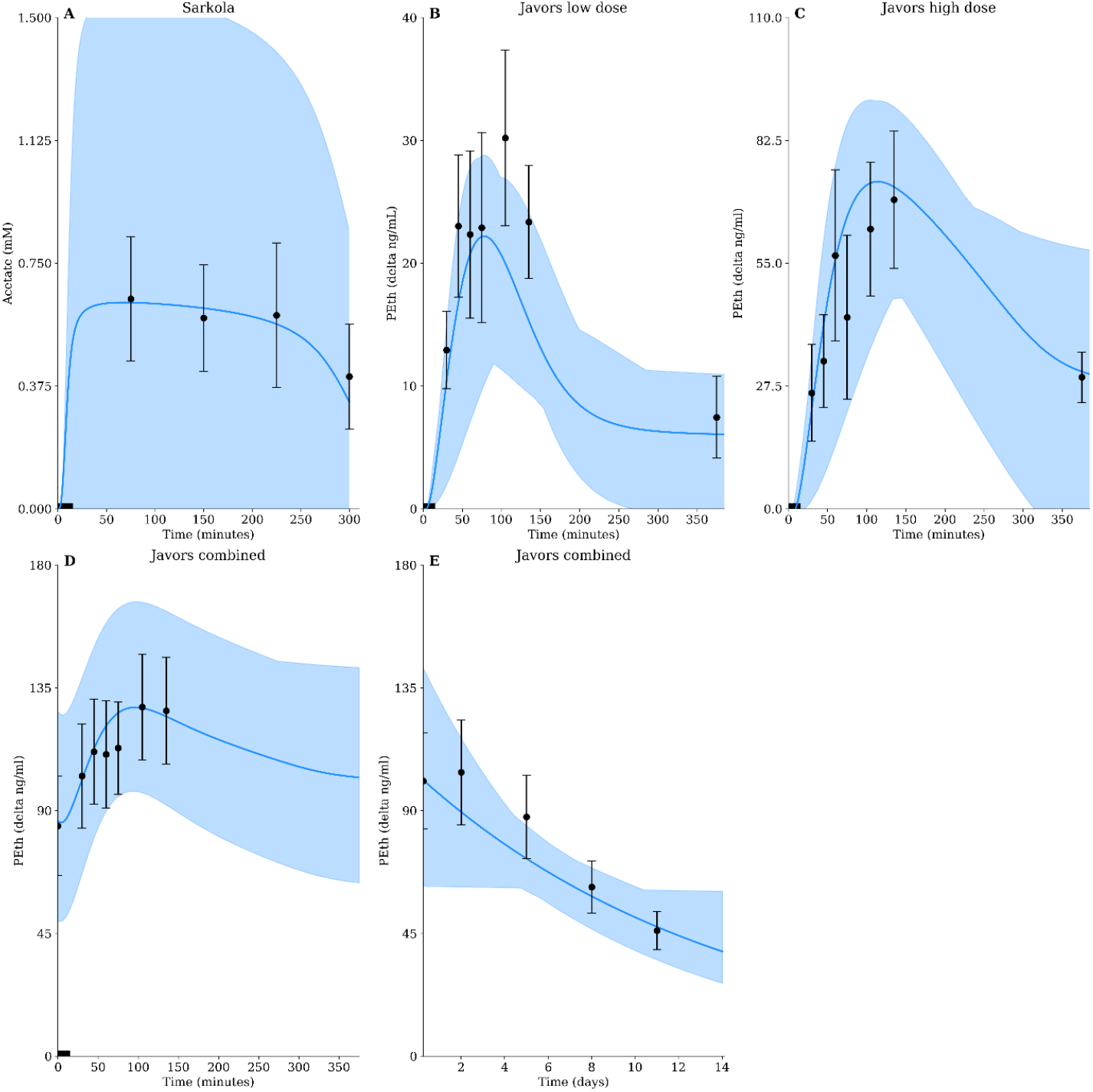
Model agreement to derivatives of oxidative and non-oxidative ethanol breakdown data for various interventions. The solid line is the best model fit, the shaded area is the model uncertainty, the black boxes are the mean experimental data values with the error bars indicating the *standard error of the mean* (SEM). The alcohol was consumed orally through different beverages. **A)** Estimated acetate from consumption of 0.28 L of 12.4 v/v % spirit blend (85 kcal) was consumed over 15 min. Estimated change of *phosphatidylethanol* (PEth), from the baseline value from consumption of **B)** 0.72L of 3.25 v/v % spirit blend (251 kcal) over 15 min and **C)** 0.71L of 6.5 v/v % spirit blend (247 kcal) over 15 min. **D)** Weighted mean behavior PEth from two groups consuming different beverages. Group 1 (n=16) consumed 0.72L of 3.25 v/v % spirit blend (251 kcal) over 15 min. Group 2 (n=11) consumed 0.71 L of 6.5 v/v % spirit blend (247 kcal) over 15 min. The combined PEth is the weighted mean behavior for both groups and shows the initial 360 minutes after the consumption. **E)** show the elimination kinetics for the following 0.25-14 days of the same experiment.

## Notes

### Competing Interest Statement

The authors have declared no competing interest.

