## Supplementary Materials for "Predicting Real-life Drinking Scenarios through a Physiological Digital Twin Incorporating Secondary Alcohol Markers"

October 21, 2025

**4 Usage of experimental data** **49**

**5 Changelog of rejected model structures** **52**

### 1 Raw prediction data

Data was collected from two individuals during a sequential drink intervention study performed by the National Board of Forensic Medicine. Written consent was obtained from all participants, and the study was approved by the Swedish Ethical Review Authority (2023-02640-01).

| **Parameter** | **BE Z00383 SD08** | **BB Z00380 SD07** |
|---|---|---|
| **Dose 1 (g/kg)** | 0.85 | 0.85 |
| **Beverage 1** | Wine (13 v/v%) | Wine (13 v/v%) |
| **Dose 2 (g/kg)** | 0.51 | 0.51 |
| **Beverage 2** | Vodka (40 v/v%) | Vodka (40 v/v%) |
| **Gender** | Male | Female |
| **Age (years)** | 28 | 22 |
| **Weight (kg)** | 106.8 | 62.7 |
| **Height (cm)** | 186 | 169 |

Supplementary Table 1: **Input values for participants in the study.** The table shows the dose of ethanol given in g/kg body weight, the beverage consumed, and participant anthropometrics.

Below the data series for the observables, BAC, UAC, EtG, and EtS, is given for the two participants.

| Time (min) | BAC (mg/dL) | UAC (mg/L) | EtG (mg/L) | EtS (mg/L) |
|---|---|---|---|---|
| 0 | 0 | 0 | 0 | 0 |
| 35 | 40.05 | - | 0 | 0.003 |
| 37 | - | 11.32 | - | - |
| 65 | 91.73 | - | 0.010 | 0.009 |
| 67 | - | 76.78 | - | - |
| 80 | 94.82 | - | 0.020 | 0.014 |
| 95 | 110.72 | - | 0.028 | 0.019 |
| 99 | - | 129.35 | - | - |
| 110 | 109.19 | - | 0.042 | 0.024 |
| 125 | 98.36 | - | 0.052 | 0.026 |
| 129 | - | 131.18 | - | - |
| 155 | 89.95 | - | 0.069 | 0.030 |
| 185 | 79.64 | - | 0.088 | 0.034 |
| 187 | - | 117.94 | - | - |
| 215 | 76.99 | - | 0.095 | 0.034 |
| 245 | 67.41 | - | 0.106 | 0.034 |
| 248 | - | 94.48 | - | - |
| 275 | 56.66 | - | 0.106 | 0.033 |
| 295 | 95.54 | - | 0.116 | 0.043 |
| 305 | 46.23 | - | 0.103 | 0.031 |
| 308 | - | 71.89 | - | - |
| 320 | 55.53 | - | 0.105 | 0.032 |
| 335 | 94.31 | - | 0.107 | 0.034 |
| 350 | 105.89 | - | 0.106 | 0.035 |
| 365 | 103.63 | - | 0.118 | 0.039 |
| 369 | - | 123.76 | - | - |
| 425 | 87.38 | - | 0.122 | 0.042 |
| 429 | - | 124.20 | - | - |
| 455 | 79.86 | - | 0.127 | 0.043 |
| 485 | 66.02 | - | 0.130 | 0.041 |
| 545 | 51.83 | 90.55 | 0.134 | 0.037 |
| 605 | 31.61 | - | 0.128 | 0.034 |
| 723 | - | 36.87 | 0.093 | 0.022 |
| 840 | 0 | - | 0.060 | 0.012 |
| 1425 | - | 0 | - | - |

Supplementary Table 2: **Experimental data for participant BE Z00383 SD08.** Missing values indicated by ”-”.

| Time (min) | BAC (mg/dL) | UAC (mg/L) | EtG (mg/L) | EtS (mg/L) |
|---|---|---|---|---|
| 0 | 0 | 0 | 0 | 0 |
| 30 | 30.01 | - | 0 | 0.002 |
| 33 | - | 11.42 | - | - |
| 60 | 76.52 | - | 0.009 | 0.010 |
| 63 | - | 90.31 | - | - |
| 75 | 82.98 | - | 0.014 | 0.014 |
| 90 | 88.03 | - | 0.021 | 0.018 |
| 92 | - | 123.10 | - | - |
| 105 | 89.00 | - | 0.032 | 0.022 |
| 120 | 87.44 | - | 0.046 | 0.028 |
| 124 | - | 115.54 | - | - |
| 135 | 90.65 | - | 0.048 | 0.027 |
| 150 | 114.01 | - | 0.060 | 0.033 |
| 165 | 126.01 | - | 0.070 | 0.038 |
| 180 | 129.53 | - | 0.077 | 0.043 |
| 185 | - | 146.57 | - | - |
| 210 | 129.70 | - | 0.103 | 0.050 |
| 240 | 117.15 | - | 0.128 | 0.055 |
| 245 | - | 159.02 | - | - |
| 270 | 109.53 | - | 0.140 | 0.055 |
| 300 | 94.52 | - | 0.145 | 0.054 |
| 305 | - | 132.96 | - | - |
| 360 | 64.63 | - | 0.158 | 0.052 |
| 364 | - | 85.88 | - | - |
| 420 | 39.72 | - | 0.131 | 0.039 |
| 425 | - | 57.13 | - | - |
| 545 | - | 11.86 | - | - |
| 600 | 29.80 | - | 0.116 | 0.030 |
| 665 | - | 0 | - | - |
| 720 | - | - | 0.075 | 0.020 |
| 1367 | - | 0 | - | - |

Supplementary Table 3: **Experimental data for participant BB Z00380 SD07.** Missing values indicated by ”-”.

### 2 Model fit to estimation data

This section present the model fit to the remaining estimation data (not included in the article) - which was also included in our previous article [1].

#### 2.1 Single drink ethanol markers

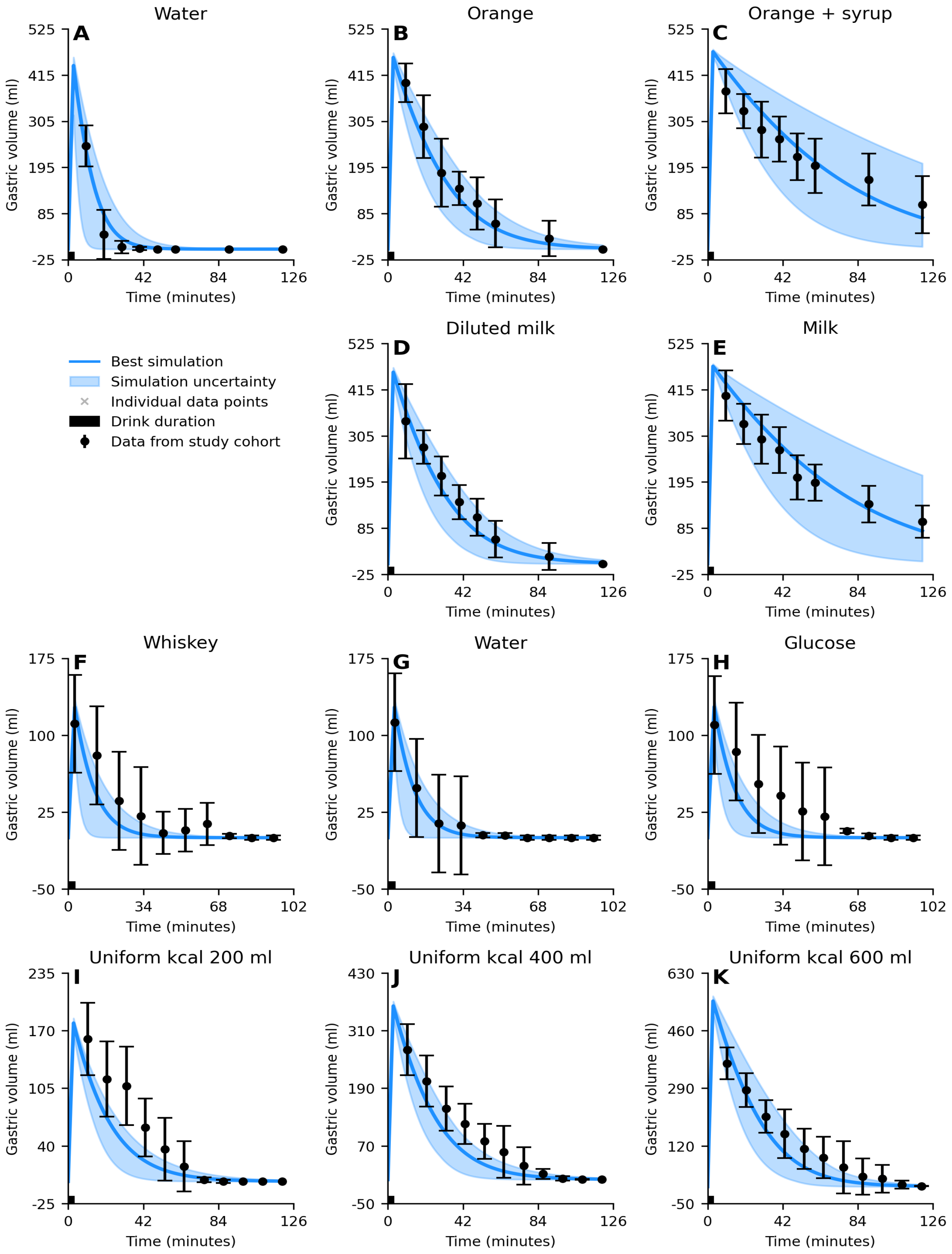

Supplementary Figure 1: *Model agreement to ethanol data for various interventions.*

**Supplementary Figure S1 (continued):** The solid line is the best model fit, the shaded area is the model uncertainty, the black boxes are the mean experimental data value with the error bars indicating the *standard error of the mean* (SEM), and individual data points are marked with a "x". The alcohol was consumed orally through different beverages in various alterations **A)** 1L of 5.1 v/v % beer (133 kcal) was consumed over 20 min, **B)** 0.42L of 12.5 v/v % wine (56 kcal) was consumed over 20 min, **C)** 0.26L of 20 v/v % spirit blend (43 kcal) was consumed over 20 min, **D)** 0.14L of 20.0 v/v % spirit blend (51 kcal) over 15 min, **E)** 0.14L of 20.0 v/v % spirit blend (51 kcal) over 15 min after a meal constituted of 700 kcal, **F)** plasma ethanol levels in women after consumption of 0.19L of 12 v/v % spirit blend (28 kcal) over 10 min after eating a meal constituted of 555 kcal. **G)** plasma ethanol levels in men after consumption of 0.22L of 12 v/v % spirit blend (33 kcal) over 10 min after eating a meal constituted of 555 kcal, **H)** plasma ethanol levels from consumption of 0.48L of 10.0 v/v % spirit blend (149 kcal) over 15 min. Estimated *blood alcohol concentration* (BAC) levels from *breath alcohol concentration* (BrAC) estimates were reported from **I)** consumption of 0.72L of 3.25 v/v % spirit blend (251 kcal) over 15 min, **J)** 0.71L of 6.5 v/v % spirit blend (247 kcal) was consumed over 15 min. In A-D and H-J the beverage was consumed in a fasting state, and in E-G in combination with food. In A-H the absolute change of BAC was measured and in I-J the absolute change in estimated BAC levels derived from BrAC measurements.

#### 2.2 Gastric emptying

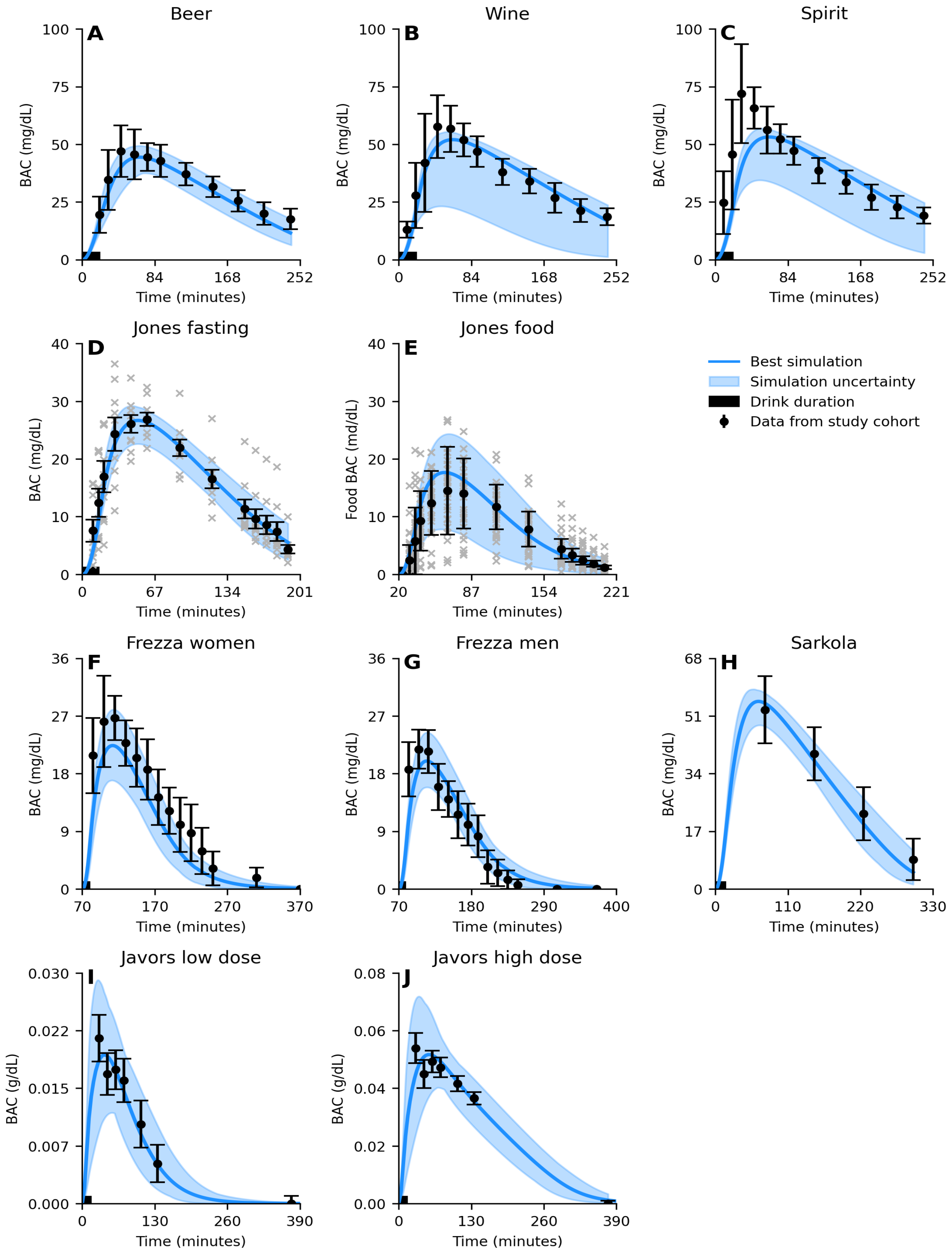

Supplementary Figure 2: *Model agreement to gastric emptying data.*

**Supplementary Figure S2 (continued):** The solid line is the best model fit, the shaded area is the model uncertainty, the black boxes are the mean experimental data value with the error bars indicating the *standard error of the mean* (SEM). The model (solid lines) describes the clearance of different beverages from the stomach (blue markers with error bars). In all experiments, the beverage was consumed over 3 minutes and was constituted of the following: **A)** 500 mL containing 0 kcal, **B)** 500 mL containing 220 kcal, **C)** 500 mL containing 329 kcal, **D)** 500 mL containing 220 kcal, **E)** 500 mL containing 329 kcal, **F)** 150 mL containing 0 non-ethanol kcal, **G)** 150 mL containing 0 kcal, **H)** 150 mL containing 67 kcal, **I)** 200 mL containing 200 kcal, **J)** 200 mL containing 400 kcal, and **K)** 600 mL containing 200 kcal.

#### 2.3 Alcohol metabolites

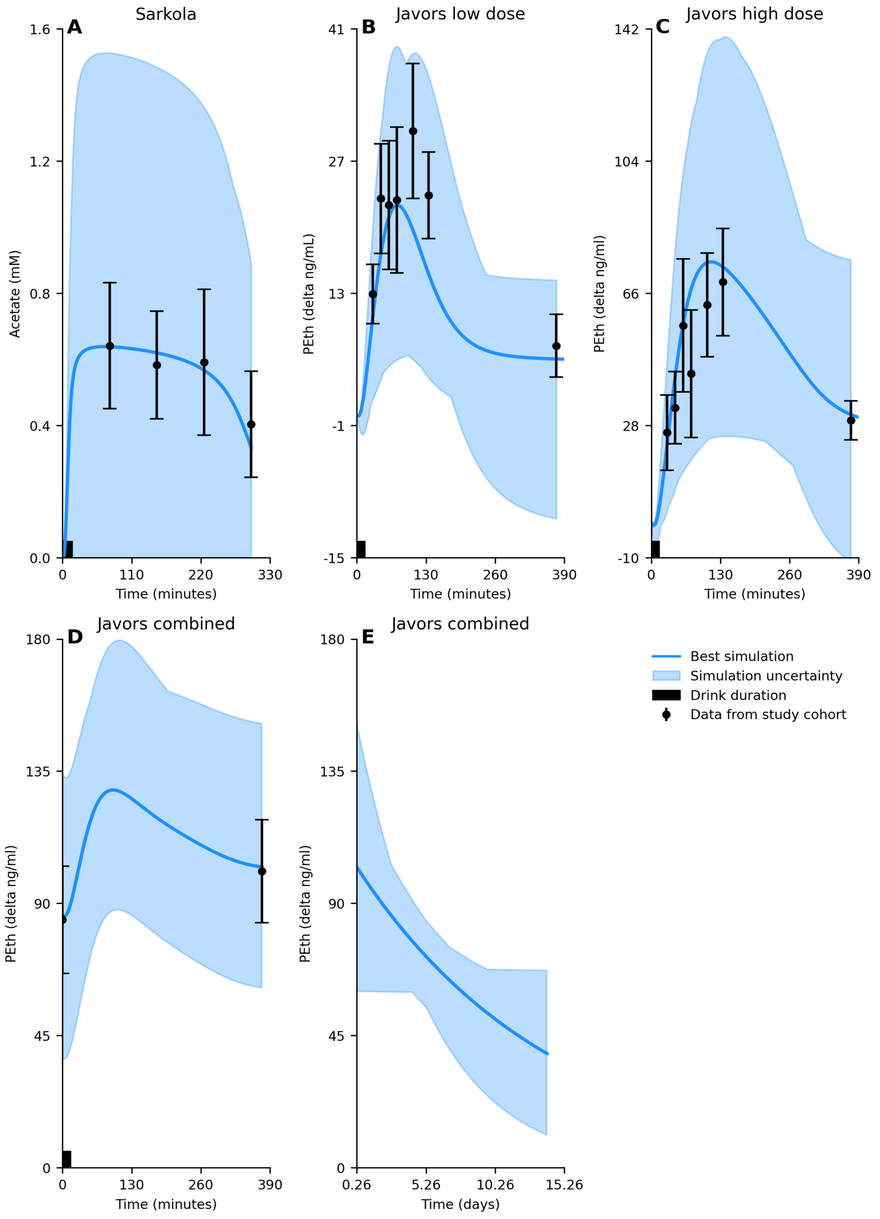

Supplementary Figure 3: *Model agreement to derivatives of oxidative and non-oxidative ethanol breakdown data for various interventions.* The solid line is the best model fit, the shaded area is the model uncertainty, the black boxes are the mean experimental data values with the error bars indicating the *standard error of the mean* (SEM). The alcohol was consumed orally through different beverages. **A)** Estimated acetate from consumption of 0.28 L of 12.4 v/v % spirit blend (85 kcal) was consumed over 15 min. Estimated change of *phosphatidylethanol* (PEth), from the baseline value from consumption of **B)** 0.72L of 3.25 v/v % spirit blend (251 kcal) over 15 min and **C)** 0.71L of 6.5 v/v % spirit blend (247 kcal) over 15 min. **D)** Weighted mean behavior PEth from two groups consuming different beverages. Group 1 (n=16) consumed 0.72L of 3.25 v/v % spirit blend (251 kcal) over 15 min. Group 2 (n=11) consumed 0.71 L of 6.5 v/v % spirit blend (247 kcal) over 15 min. The combined PEth is the weighted mean behavior for both groups and shows the initial 360 minutes after the consumption. **E)** show the elimination kinetics for the following 0.25-14 days of the same experiment.

### 3 Input estimations

This section presents all the inferred values that has been estimated for the different data sets and how this was performed.

#### 3.1 Methods for estimating values

##### 3.1.1 Estimating the height of the participants when no height where given

We used databases to estimate the height, using the national averages in from the country of origin of the original paper, and if possible also the national average for the year the paper was published. All such values are given in the tables below.

##### 3.1.2 Calculating weight from BMI

If the BMI of the participants were given in the original paper but not the weight, then we calculated the weight based on the height of the participants and the BMI, using the definition of BMI.

$$weight = BMI \cdot height^2 \tag{1}$$

##### 3.1.3 Estimating drink volume from dose and concentration

In many studies, the full details of the drinks were not given in the paper. The dose of alcohol given was often defined as g ethanol per kg body weight of the participants. Thus we could calculate the total grams of alcohol given by multiplying the ethanol g/kg body weight (X) with the body weight:

$$mass_{ethanol} = X \cdot weight \tag{2}$$

Using the density of alcohol (g/ml), we could calculate the total volume (ml) of ethanol in the drink:

$$V_{ethanol} = \frac{mass_{ethanol}}{0.7891} \tag{3}$$

Knowing the final volume of ethanol in the drink, we could calculate the total volume using the dilution equation:

$$c_{ethanol} \cdot V_{ethanol} = c_{drink} \cdot V_{drink} \tag{4}$$

Here we assume that the concentration of ethanol is 100%, thus $c_{ethanol} = 1$ and:

$$V_{drink} = \frac{vol_{ethanol}}{c_{drink}} \tag{5}$$

##### 3.1.4 Average sex in a group

In some datasets the participants were both men and women. From the perspective of the model, the major difference between individual were driven by the differences in total blood volume. Thus, if the dataset contained both men and women, we needed to construct an average person (either man or woman) but with the average blood volume. For this, we first determined if men or women were the most common sex in the group, and then we calculated the blood volumes for the subsection of either sex, and then constructed an average individual. We calculated the blood volume using the Nadler equation [2]. For men blood volume was estimated using, where the height is given in meters:

$$vol_{blood,male} = 0.3669 \cdot height_{male}^3 + 0.03219 \cdot weight_{male} + 0.6041 \tag{6}$$

And for women:

$$vol_{blood,female} = 0.3561 \cdot height_{female}^3 + 0.03308 \cdot weight_{female} + 0.1833) \tag{7}$$

We then calculated the ratio of women to men according to, where sex=1 if man, and sex=0 if woman:

$$sex = \frac{n_{males}}{n_{females} + n_{males}} \tag{8}$$

We then calculated the average blood volume within the group:

$$vol_{blood} = sex \cdot vol_{blood,male} + (1 - sex) \cdot vol_{blood,female} \tag{9}$$

From the average blood volume, we reverse calculated an individual with the same blood volume. If there were more men in the group we assumed the height to be the average height within the men in the group, and then calculated the weight using the Nadler equation according to:

$$height_{male} = \left( \frac{vol_{blood} - 0.6041 - 0.03219 \cdot weight}{0.3669} \right)^{\frac{1}{3}} \tag{10}$$

If there were more women than men, we instead used the average weight of the women, and calculated the height according to:

$$height_{female} = \left( \frac{vol_{blood} - 0.1833 - 0.03308 \cdot weight}{0.3561} \right)^{\frac{1}{3}} \tag{11}$$

###### 3.1.5 Estimating calories within the drink

If the authors of the original papers had not disclosed the caloric content of the drinks, we approximated the calories based on the drink type. Note that the calories of the drink excludes the calories from the ethanol.

#### 3.2 Information for each dataset

Below are the values used for the different experiments. If the value were given in the paper, the value in the "Given values" column was used, and if not, we inferred the value as described above and used the value in the "Inferred values" column. The different inputs correspond to:

- Sex: the most frequent sex of the participants. 0 corresponds to women, and 1 corresponds to men.
- Height: the height in meters
- Weight: the weight in kg
- Age: the age in years
- BMI: the body mass index. Was used to infer the weight if no information on the weight was given.

- Volume: The volume of the drink consumed
- Concentration: The concentration of the drink consumed (in v/v%)
- Drink kcal: the amount of kilocalories in the drink, excluding the calories from ethanol
- Drink length: under how long time (minutes) the drink was consumed
- Food kcal: the amount of kilocalories in food consumed
- Time of meal: the time when the meal was consumed (relative to the start of the drink)

Additionally, time of urination was given as for the Kronstrand *et al.* [3] and Hoiseth *et al.* [4] for the following time values: 30.01, 60.01, 90.01, 120.01, 180.01, 240.01, 300.01, 360.01, 420.01.
Below we give the values for each dataset/experiment.

##### 3.2.1 Kronstrand low dose whiskey

| **Input** | **Given values** | **Inferred values** |
|---|---|---|
| Sex | 1 | – |
| Height | 1.71 | – |
| Weight | – | 79.94 |
| Age | 29 | – |
| BMI | – | 27.34 |
| Volume | – | 4 · 0.258, 0.127 |
| Concentration | 4 · 5.2 | 20 |
| Drink kcal | – | 4 · 33.54, 25.33 |
| Drink length | 4 · 10, 15 | – |
| Food kcal | – | 300, 500 |
| Time of meal | – | 0, 180 |

Supplementary Table 4: **Input values for Kronstrand low dose whiskey**

###### 3.2.2 Kronstrand medium dose whiskey

| Input | Given values | Inferred values |
|---|---|---|
| Sex | 1 | – |
| Height | 1.74 | – |
| Weight | – | 78.18 |
| Age | 34 | – |
| BMI | – | 24.67 |
| Volume | – | $4 \cdot 0.253$, 0.253 |
| Concentration | $4 \cdot 5.2$ | 20 |
| Drink kcal | – | $4 \cdot 32.80$, 50.53 |
| Drink length | $4 \cdot 10$, 30 | – |
| Food kcal | – | 300, 500 |
| Time of meal | – | 0, 180 |

Supplementary Table 5: **Input values for Kronstrand medium dose whiskey**

###### 3.2.3 Kronstrand high dose whiskey

| Input | Given values | Inferred values |
|---|---|---|
| Sex | 1 | – |
| Height | 1.73 | – |
| Weight | – | 67.20 |
| Age | 28 | – |
| BMI | – | 22.45 |
| Volume | – | $4 \cdot 0.217$, 0.362 |
| Concentration | $4 \cdot 5.2$ | 20 |
| Drink kcal | – | $4 \cdot 28.19$, 72.40 |
| Drink length | $4 \cdot 10$, 30 | – |
| Food kcal | – | 300, 500 |
| Time of meal | – | 0, 180 |

Supplementary Table 6: **Input values for Kronstrand high dose whiskey**

###### 3.2.4 Hoiseth low dose beer

| Input | Given values | Inferred values |
|---|---|---|
| Sex | 1 | – |
| Height | 1.66 | – |
| Weight | – | 70.70 |
| Age | 22 | – |
| BMI | – | 25.66 |
| Volume | – | $4 \cdot 0.204$, 0.40 |
| Concentration | $4 \cdot 5.6$, 5.6 | – |
| Drink kcal | – | $4 \cdot 26.49$, 51.93 |
| Drink length | $4 \cdot 10$, 15 | – |
| Food kcal | – | 300, 500 |
| Time of meal | – | 0, 180 |

Supplementary Table 7: **Input values for Hoiseth low dose beer**

###### 3.2.5 Hoiseth medium dose beer

| Input | Given values | Inferred values |
|---|---|---|
| Sex | 1 | – |
| Height | 1.81 | – |
| Weight | – | 79.77 |
| Age | 24 | – |
| BMI | – | 24.35 |
| Volume | – | $4 \cdot 0.230$, 0.921 |
| Concentration | $4 \cdot 5.6$, 5.6 | – |
| Drink kcal | – | $4 \cdot 29.88$, 119.53 |
| Drink length | $4 \cdot 10$, 30 | – |
| Food kcal | – | 300, 500 |
| Time of meal | – | 0, 180 |

Supplementary Table 8: **Input values for Hoiseth medium dose beer**

##### 3.2.6 Hoiseth low dose wine

| Input | Given values | Inferred values |
|---|---|---|
| Sex | 1 | – |
| Height | 1.79 | – |
| Weight | – | 76.62 |
| Age | 22 | – |
| BMI | – | 23.91 |
| Volume | – | $4 \cdot 0.221$, 0.187 |
| Concentration | $4 \cdot 5.6$, 13 | – |
| Drink kcal | – | $4 \cdot 28.70$, 24.88 |
| Drink length | $4 \cdot 10$, 15 | – |
| Food kcal | – | 300, 500 |
| Time of meal | – | 0, 180 |

Supplementary Table 9: **Input values for Hoiseth low dose wine**

##### 3.2.7 Hoiseth medium dose wine

| Input | Given values | Inferred values |
|---|---|---|
| Sex | 1 | – |
| Height | 1.79 | – |
| Weight | – | 73.54 |
| Age | 23 | – |
| BMI | – | 22.95 |
| Volume | – | $4 \cdot 0.212$, 0.366 |
| Concentration | $4 \cdot 5.6$, 13 | – |
| Drink kcal | – | $4 \cdot 27.55$, 48.72 |
| Drink length | $4 \cdot 10$, 30 | – |
| Food kcal | – | 300, 500 |
| Time of meal | – | 0, 180 |

Supplementary Table 10: **Input values for Hoiseth medium dose wine**

###### 3.2.8 Hoiseth low dose vodka

| Input | Given values | Inferred values |
|---|---|---|
| Sex | 0 | – |
| Height | 1.72 | – |
| Weight | – | 69.84 |
| Age | 24 | – |
| BMI | – | 23.61 |
| Volume | – | $4 \cdot 0.202$, 0.111 |
| Concentration | $4 \cdot 5.6$ | 20 |
| Drink kcal | – | $4 \cdot 26.16$, 22.13 |
| Drink length | $4 \cdot 10$, 15 | – |
| Food kcal | – | 300, 500 |
| Time of meal | – | 0, 180 |

Supplementary Table 11: **Input values for Hoiseth low dose vodka**

###### 3.2.9 Hoiseth medium dose vodka

| Input | Given values | Inferred values |
|---|---|---|
| Sex | 1 | – |
| Height | 1.78 | – |
| Weight | – | 82.18 |
| Age | 23 | – |
| BMI | – | 25.94 |
| Volume | – | $4 \cdot 0.237$, 0.266 |
| Concentration | $4 \cdot 5.6$ | 20 |
| Drink kcal | – | $4 \cdot 30.79$, 53.12 |
| Drink length | $4 \cdot 10$, 30 | – |
| Food kcal | – | 300, 500 |
| Time of meal | – | 0, 180 |

Supplementary Table 12: **Input values for Hoiseth medium dose vodka**

###### 3.2.10 Hoiseth high dose vodka

| Input | Given values | Inferred values |
|---|---|---|
| Sex | 1 | – |
| Height | 1.82 | – |
| Weight | – | 80.78 |
| Age | 23 | – |
| BMI | – | 24.39 |
| Volume | – | $4 \cdot 0.233$, 0.435 |
| Concentration | $4 \cdot 5.6$ | 20 |
| Drink kcal | – | $4 \cdot 30.26$, 87.03 |
| Drink length | $4 \cdot 10$, 20 | – |
| Food kcal | – | 300, 500 |
| Time of meal | – | 0, 180 |

Supplementary Table 13: **Input values for Hoiseth high dose vodka**

###### 3.2.11 Wang

| Input | Given values | Inferred values |
|---|---|---|
| Sex | 1 | – |
| Height | – | 1.66 |
| Weight | – | 52.2 |
| Age | 24.5 | – |
| BMI | 20.9 | – |
| Volume | – | 0.119 |
| Concentration | 40 | – |
| Drink kcal | 0 | – |
| Drink length | 30 | – |
| Food kcal | 500 | – |
| Time of meal | 0 | – |

Supplementary Table 14: **Input values for Wang**

##### 3.2.12 Okabe 500 ml Water

| **Input** | **Given values** | **Inferred values** |
|---|---|---|
| Sex | 1 | – |
| Height | 1.71 | – |
| Weight | 65 | – |
| Age | 27 | – |
| BMI | 22 | – |
| Volume | 0.5 | – |
| Concentration | 0.0 | – |
| Drink kcal | 0 | – |
| Drink length | 3 | – |
| Food kcal | 0 | – |
| Time of meal | 0 | – |

Supplementary Table 15: **Input values for Okabe 500 ml water**

##### 3.2.13 Okabe 500 ml orange juice

| **Input** | **Given values** | **Inferred values** |
|---|---|---|
| Sex | 1 | – |
| Height | 1.71 | – |
| Weight | 65 | – |
| Age | 27 | – |
| BMI | 22 | – |
| Volume | 0.5 | – |
| Concentration | 0.0 | – |
| Drink kcal | 220 | – |
| Drink length | 3 | – |
| Food kcal | 0 | – |
| Time of meal | 0 | – |
| Time of urination | – | – |

Supplementary Table 16: **Input values for Okabe 500 ml orange juice**

###### 3.2.14 Okabe 500 ml milk water

| Input | Given values | Inferred values |
|---|---|---|
| Sex | 1 | – |
| Height | 1.71 | – |
| Weight | 65 | – |
| Age | 27 | – |
| BMI | 22 | – |
| Volume | 0.5 | – |
| Concentration | 0.0 | – |
| Drink kcal | 220 | – |
| Drink length | 3 | – |
| Food kcal | 0 | – |
| Time of meal | 0 | – |

Supplementary Table 17: **Input values for Okabe 500 ml milk water**

###### 3.2.15 Okabe 500 ml orange juice and syrup

| Input | Given values | Inferred values |
|---|---|---|
| Sex | 1 | – |
| Height | 1.71 | – |
| Weight | 65 | – |
| Age | 27 | – |
| BMI | 22 | – |
| Volume | 0.5 | – |
| Concentration | 0.0 | – |
| Drink kcal | 330 | – |
| Drink length | 3 | – |
| Food kcal | 0 | – |
| Time of meal | 0 | – |

Supplementary Table 18: **Input values for Okabe 500 ml orange juice and syrup**

###### 3.2.16 Okabe 500 ml milk

| Input | Given values | Inferred values |
|---|---|---|
| Sex | 1 | – |
| Height | 1.71 | – |
| Weight | 65 | – |
| Age | 27 | – |
| BMI | 22 | – |
| Volume | 0.5 | – |
| Concentration | 0.0 | – |
| Drink kcal | 330 | – |
| Drink length | 3 | – |
| Food kcal | 0 | – |
| Time of meal | 0 | – |

Supplementary Table 19: **Input values for Okabe 500 ml milk**

###### 3.2.17 Okabe 150 ml water

| Input | Given values | Inferred values |
|---|---|---|
| Sex | 1 | – |
| Height | 1.712 | – |
| Weight | 68.4 | – |
| Age | 32 | – |
| BMI | 23 | – |
| Volume | 0.15 | – |
| Concentration | 0.0 | – |
| Drink kcal | 0 | – |
| Drink length | 3 | – |
| Food kcal | 0 | – |
| Time of meal | 0 | – |

Supplementary Table 20: **Input values for Okabe 150 ml water**

###### 3.2.18 Okabe 150 ml glucose

| **Input** | **Given values** | **Inferred values** |
|---|---|---|
| Sex | 1 | – |
| Height | 1.712 | – |
| Weight | 68.4 | – |
| Age | 32 | – |
| BMI | 23 | – |
| Volume | 0.15 | – |
| Concentration | 0.0 | – |
| Drink kcal | 67 | – |
| Drink length | 3 | – |
| Food kcal | 0 | – |
| Time of meal | 0 | – |
| Time of urination | – | – |

Supplementary Table 21: **Input values for Okabe 150 ml glucose**

###### 3.2.19 Okabe 150 ml whiskey

| **Input** | **Given values** | **Inferred values** |
|---|---|---|
| Sex | 1 | – |
| Height | 1.712 | – |
| Weight | 68.4 | – |
| Age | 32 | – |
| BMI | 23 | – |
| Volume | 0.15 | – |
| Concentration | 0.0 | – |
| Drink kcal | 0 | – |
| Drink length | 3 | – |
| Food kcal | 0 | – |
| Time of meal | 0 | – |

Supplementary Table 22: **Input values for Okabe 150 ml whiskey**

###### 3.2.20 Okabe 200 ml uniform glucose

| Input | Given values | Inferred values |
|---|---|---|
| Sex | 1 | – |
| Height | 1.69 | – |
| Weight | 60.0 | – |
| Age | 27 | – |
| BMI | 21 | – |
| Volume | 0.2 | – |
| Concentration | 0.0 | – |
| Drink kcal | 200 | – |
| Drink length | 3 | – |
| Food kcal | 0 | – |
| Time of meal | 0 | – |

Supplementary Table 23: **Input values for Okabe 200 ml uniform glucose**

###### 3.2.21 Okabe 400 ml uniform glucose

| Input | Given values | Inferred values |
|---|---|---|
| Sex | 1 | – |
| Height | 1.69 | – |
| Weight | 60.0 | – |
| Age | 27 | – |
| BMI | 21 | – |
| Volume | 0.4 | – |
| Concentration | 0.0 | – |
| Drink kcal | 200 | – |
| Drink length | 3 | – |
| Food kcal | 0 | – |
| Time of meal | 0 | – |

Supplementary Table 24: **Input values for Okabe 400 ml uniform glucose**

###### 3.2.22 Okabe 600 ml uniform glucose

| Input | Given values | Inferred values |
|---|---|---|
| Sex | 1 | – |
| Height | 1.69 | – |
| Weight | 60.0 | – |
| Age | 27 | – |
| BMI | 21 | – |
| Volume | 0.6 | – |
| Concentration | 0.0 | – |
| Drink kcal | 200 | – |
| Drink length | 3 | – |
| Food kcal | 0 | – |
| Time of meal | 0 | – |

Supplementary Table 25: **Input values for Okabe 600 ml uniform glucose**

###### 3.2.23 Mitchell beer

| Input | Given values | Inferred values |
|---|---|---|
| Sex | 1 | – |
| Height | – | 1.77 |
| Weight | 82.66 | – |
| Age | 25 | – |
| BMI | 26.35 | – |
| Volume | – | 1.027 |
| Concentration | 5.1 | – |
| Drink kcal | – | 133.33 |
| Drink length | 20 | – |
| Food kcal | 0 | – |
| Time of meal | 0 | – |

Supplementary Table 26: **Input values for Mitchell beer**

###### 3.2.24 Mitchell wine

| Input | Given values | Inferred values |
|---|---|---|
| Sex | 1 | – |
| Height | – | 1.77 |
| Weight | 82.66 | – |
| Age | 25 | – |
| BMI | 26.35 | – |
| Volume | – | 0.419 |
| Concentration | 12.5 | – |
| Drink kcal | – | 55.83 |
| Drink length | 20 | – |
| Food kcal | 0 | – |
| Time of meal | 0 | – |

Supplementary Table 27: **Input values for Mitchell wine**

###### 3.2.25 Mitchell spirit

| Input | Given values | Inferred values |
|---|---|---|
| Sex | 1 | – |
| Height | – | 1.77 |
| Weight | 82.66 | – |
| Age | 85 | – |
| BMI | 26.35 | – |
| Volume | – | 0.2618 |
| Concentration | 20.0 | – |
| Drink kcal | – | 43.18 |
| Drink length | 20 | – |
| Food kcal | 0 | – |
| Time of meal | 0 | – |

Supplementary Table 28: **Input values for Mitchell spirit**

###### 3.2.26 Jones fasting

| Input | Given values | Inferred values |
|---|---|---|
| Sex | 1 | – |
| Height | 1.83 | – |
| Weight | 75.8 | – |
| Age | 25 | – |
| BMI | – | 22.63 |
| Volume | – | 0.1441 |
| Concentration | 20 | 9.44 |
| Drink kcal | – | 51.877 |
| Drink length | 15 | – |
| Food kcal | 0 | – |
| Time of meal | 0 | – |

Supplementary Table 29: **Input values for Jones fasting**

###### 3.2.27 Jones food

| Input | Given values | Inferred values |
|---|---|---|
| Sex | 1 | – |
| Height | 1.83 | – |
| Weight | 75.8 | – |
| Age | 25 | – |
| BMI | – | 22.63 |
| Volume | – | 0.1441 |
| Concentration | 20 | 9.44 |
| Drink kcal | – | 51.877 |
| Drink length | 15 | – |
| Food kcal | 700 | – |
| Time of meal | -20 | – |

Supplementary Table 30: **Input values for Jones food**

###### 3.2.28 Sarkola

| Input | Given values | Inferred values |
|---|---|---|
| Sex | – | 0 |
| Height | – | 1.762 |
| Weight | – | 65.30 |
| Age | 21 | – |
| BMI | – | 21.03 |
| Volume | – | 0.414 |
| Concentration | 10 | – |
| Drink kcal | – | 133.33 |
| Drink length | 15 | – |
| Food kcal | 0 | – |
| Time of meal | 0 | – |

Supplementary Table 31: **Input values for Sarkola**

###### 3.2.29 Javors low dose

| Input | Given values | Inferred values |
|---|---|---|
| Sex | – | 1 |
| Height | – | 1.78 |
| Weight | 58.92 | – |
| Age | 28 | – |
| BMI | – | 18.60 |
| Volume | 0.70976 | – |
| Concentration | – | 3.245 |
| Drink kcal | – | 242.05 |
| Drink length | 15 | – |
| Food kcal | 0 | – |
| Time of meal | 0 | – |

Supplementary Table 32: **Input values for Javors low dose**

###### 3.2.30 Javors high dose

| Input | Given values | Inferred values |
|---|---|---|
| Sex | – | 1 |
| Height | – | 1.78 |
| Weight | 59.23 | – |
| Age | 28 | – |
| BMI | – | 18.70 |
| Volume | 0.70976 | – |
| Concentration | – | 6.41 |
| Drink kcal | – | 235.85 |
| Drink length | 15 | – |
| Food kcal | 0 | – |
| Time of meal | 0 | – |

Supplementary Table 33: **Input values for Javors high dose**

###### 3.2.31 Frezza woman

| Input | Given values | Inferred values |
|---|---|---|
| Sex | 0 | – |
| Height | – | 1.61 |
| Weight | – | 59.62 |
| Age | – | 35 |
| BMI | – | 23 |
| Volume | – | 0.1889 |
| Concentration | – | 12 |
| Drink kcal | – | 28.25 |
| Drink length | 10 | – |
| Food kcal | 555 | – |
| Time of meal | -60 | – |

Supplementary Table 34: **Input values for Frezza woman**

###### 3.2.32 Frezza men

| Input | Given values | Inferred values |
|---|---|---|
| Sex | 1 | – |
| Height | – | 1.74 |
| Weight | – | 69.63 |
| Age | – | 40 |
| BMI | – | 23 |
| Volume | – | 0.221 |
| Concentration | – | 12 |
| Drink kcal | – | 33 |
| Drink length | 10 | – |
| Food kcal | 555 | – |
| Time of meal | -60 | – |
| Time of urination | -/ | – |

Supplementary Table 35: **Input values for Frezza men**

#### 3.3 Calculations for inferred values

##### 3.3.1 Kronstrand low dose whiskey

**Calculation of a mean person**

The blood volume was calculated for the individuals in the group, using Nadler's equation [2], and then the group height was inferred from Nadler's equation - using the mean blood volume, mean sex, and mean weight.

$$
\begin{aligned}
&n_{females} = 1\\
&n_{males} = 4\\
&sex = \frac{n_{males}}{n_{females} + n_{males}} = \frac{4}{1+4} = 0.8\\
&mean_{sex} = 1\\
&mean_{weight} = 74.94 \text{ kg}\\
&mean_{bloodVolume} = 48.4833 \text{ dL}\\
&height = \left(\frac{mean_{bloodVolume} - 0.03219 \cdot mean_{weight} + 0.6041}{0.3669}\right)^{\frac{1}{3}}\\
&height = \left(\frac{48.4833 - 0.03219 \cdot 74.94 + 0.6041}{0.3669}\right)^{\frac{1}{3}}\\
&height = 1.709 \text{ m}
\end{aligned}
\tag{12}
$$

**Estimated inputs**

Knowing that the density of ethanol is equal to 0.7891 g/ml or kg/L and the reported consumption of 0.51 g ethanol per kg bodyweight beer and 0.25 g per kg bodyweight whiskey [3], we can estimate the inputs. We assume that the 1h period of the beer was equally consumed over 4 blocks of 10 minutes to account for the blood and urine samples that was taken. We also assume that the subjects drank equal volume of non-alcoholic beverages alongside the spirits - effectively doubling the volume and halving the concentration (20 v/v%).

$$
\begin{aligned}
mass_{EtOHBeer} &= weight \cdot 0.51 = 74.94 \cdot 0.51 = 38.219 \text{ g} \\
volume_{EtOHBeer} &= \frac{38.219}{0.7891 \cdot 1000} = 0.0484 \text{ L} \\
volume_{beer} &= \frac{c2 \cdot v2}{c1} = \frac{0.0484 \cdot 1}{0.052} = 0.931 \text{ L} \\
volume_{beerQuarter} &= \frac{0.931}{4} = 0.233 \text{ L} \\
mass_{EtOHWhiskey} &= weight \cdot 0.25 = 74.94 \cdot 0.25 = 18.735 \text{ g} \\
volume_{EtOHWhiskey} &= \frac{18.735}{0.7891 \cdot 1000} = 0.0237 \text{ L} \\
volume_{whiskey} &= \frac{c2 \cdot v2}{c1} = \frac{0.0237 \cdot 1}{0.20} = 0.1185 \text{ L}
\end{aligned}
\tag{13}
$$

##### 3.3.2 Kronstrand medium dose whiskey

###### Calculation of a mean person

The blood volume was calculated for the individuals in the group, using Nadler's equation [2], and then the group height was inferred from Nadler's equation - using the mean blood volume, mean sex, and mean weight.

$$
\begin{aligned}
&n_{females} = 1 \\
&n_{males} = 4 \\
&sex = \frac{n_{males}}{n_{females} + n_{males}} = \frac{4}{1+4} = 0.8 \\
&mean_{sex} = 1 \\
&mean_{weight} = 78.18 \text{ kg} \\
&mean_{bloodVolume} = 50.5194 \text{ dL} \\
&height = \left(\frac{mean_{bloodVolume} - 0.03219 \cdot mean_{weight} + 0.6041}{0.3669}\right)^{\frac{1}{3}} \\
&height = \left(\frac{50.5194 - 0.03219 \cdot 78.18 + 0.6041}{0.3669}\right)^{\frac{1}{3}} \\
&height = 1.739 \text{ m}
\end{aligned}
\tag{14}
$$

###### Estimated inputs

Knowing that the density of ethanol is equal to 0.7891 g/ml or kg/L and the reported consumption of 0.51 g ethanol per kg bodyweight beer and 0.51 g per kg bodyweight whiskey [3], we can estimate the inputs. We assume that the 1h period of the beer was equally consumed over 4 blocks of 10 minutes to account for the blood and urine samples that was taken. We also assume that the subjects drank equal volume of non-alcoholic beverages alongside the spirits - effectively doubling the volume and halving the concentration (20 v/v%).

$$
\begin{aligned}
&mass_{EtOHBeer} = weight \cdot 0.51 = 78.18 \cdot 0.51 = 39.872 \text{ g} \\
&volume_{EtOHBeer} = \frac{39.872}{0.7891 \cdot 1000} = 0.0505 \text{ L} \\
&volume_{beer} = \frac{c2 \cdot v2}{c1} = \frac{0.0505 \cdot 1}{0.052} = 0.971 \text{ L} \\
&volume_{beerQuarter} = \frac{0.971}{4} = 0.243 \text{ L} \\
&mass_{EtOHWhiskey} = weight \cdot 0.51 = 78.18 \cdot 0.51 = 39.872 \text{ g} \\
&volume_{EtOHWhiskey} = \frac{39.872}{0.7891 \cdot 1000} = 0.0505 \text{ L} \\
&volume_{whiskey} = \frac{c2 \cdot v2}{c1} = \frac{0.0505 \cdot 1}{0.20} = 0.2525 \text{ L}
\end{aligned} \tag{15}
$$

##### 3.3.3 Kronstrand high dose whiskey

###### Calculation of a mean person

The blood volume was calculated for the individuals in the group, using Nadler's equation [2], and then the group height was inferred from Nadler's equation - using the mean blood volume, mean sex, and mean weight.

$$
\begin{aligned}
&n_{females} = 2 \\
&n_{males} = 3 \\
&sex = \frac{n_{males}}{n_{females} + n_{males}} = \frac{3}{2+3} = 0.6 \\
&mean_{sex} = 1 \\
&mean_{weight} = 67.18 \text{ kg} \\
&mean_{bloodVolume} = 46.5852 \text{ dL} \\
&height = \left( \frac{mean_{bloodVolume} - 0.03219 \cdot mean_{weight} + 0.6041}{0.3669} \right)^{\frac{1}{3}} \\
&height = \left( \frac{46.5852 - 0.03219 \cdot 67.18 + 0.6041}{0.3669} \right)^{\frac{1}{3}} \\
&height = 1.728 \text{ m}
\end{aligned} \tag{16}
$$

###### Estimated inputs

Knowing that the density of ethanol is equal to 0.7891 g/ml or kg/L and the reported consumption of 0.51 g ethanol per kg bodyweight beer and 0.85 g per kg bodyweight whiskey [3], we can estimate the inputs. We assume that the 1h period of the beer was equally consumed over 4 blocks of 10 minutes to account for the blood and urine samples that was taken. We also assume that the subjects drank equal volume of non-alcoholic beverages alongside the spirits - effectively doubling the volume and halving the concentration (20 v/v%).

$$
\begin{aligned}
&mass_{EtOHBeer} = weight \cdot 0.51 = 67.18 \cdot 0.51 = 36.262 \text{ g} \\
&volume_{EtOHBeer} = \frac{36.262}{0.7891 \cdot 1000} = 0.046 \text{ L} \\
&volume_{beer} = \frac{c2 \cdot v2}{c1} = \frac{0.046 \cdot 1}{0.052} = 0.885 \text{ L} \\
&volume_{beerQuarter} = \frac{0.885}{4} = 0.221 \text{ L} \\
&mass_{EtOHWhiskey} = weight \cdot 0.85 = 67.18 \cdot 0.85 = 57.103 \text{ g} \\
&volume_{EtOHWhiskey} = \frac{57.103}{0.7891 \cdot 1000} = 0.072 \text{ L} \\
&volume_{whiskey} = \frac{c2 \cdot v2}{c1} = \frac{0.072 \cdot 1}{0.20} = 0.36 \text{ L}
\end{aligned}
\tag{17}
$$

##### 3.3.4 Hoiseth low dose beer

#### Calculation of a mean person

The blood volume was calculated for the individuals in the group, using Nadler's equation [2], and then the group height was inferred from Nadler's equation - using the mean blood volume, mean sex, and mean weight.

$$
\begin{aligned}
&n_{females} = 2 \\
&n_{males} = 2 \\
&sex = \frac{n_{males}}{n_{females} + n_{males}} = \frac{2}{2+2} = 0.5 \\
&mean_{sex} = 1 \\
&mean_{weight} = 70.70 \text{ kg} \\
&mean_{bloodVolume} = 45.7176 \text{ dL} \\
&height = \left( \frac{mean_{bloodVolume} - 0.03219 \cdot mean_{weight} + 0.6041}{0.3669} \right)^{\frac{1}{3}} \\
&height = \left( \frac{45.7176 - 0.03219 \cdot 70.70 + 0.6041}{0.3669} \right)^{\frac{1}{3}} \\
&height = 1.664 \text{ m}
\end{aligned}
\tag{18}
$$

#### Estimated inputs

Knowing that the density of ethanol is equal to 0.7891 g/ml or kg/L and the reported consumption of 0.51 g ethanol per kg bodyweight beer and 0.25 g per kg bodyweight beer [4], we can estimate the inputs. We assume that the 1h period of the beer was equally consumed over 4 blocks of 10 minutes to account for the blood and urine samples that was taken.

$$
\begin{aligned}
&mass_{EtOHBeer} = weight \cdot 0.51 = 70.70 \cdot 0.51 = 36.057 \text{ g} \\
&volume_{EtOHBeer} = \frac{36.057}{0.7891 \cdot 1000} = 0.0457 \text{ L} \\
&volume_{beer} = \frac{c2 \cdot v2}{c1} = \frac{0.0457 \cdot 1}{0.056} = 0.816 \text{ L} \\
&volume_{beerQuarter} = \frac{0.931}{4} = 0.204 \text{ L} \\
&mass_{EtOHWhiskey} = weight \cdot 0.25 = 70.70 \cdot 0.25 = 17.675 \text{ g} \\
&volume_{EtOHWhiskey} = \frac{17.675}{0.7891 \cdot 1000} = 0.0224 \text{ L} \\
&volume_{whiskey} = \frac{c2 \cdot v2}{c1} = \frac{0.0224 \cdot 1}{0.056} = 0.4 \text{ L}
\end{aligned} \tag{19}
$$

##### 3.3.5 Hoiseth medium dose beer

#### Calculation of a mean person

The blood volume was calculated for the individuals in the group, using Nadler's equation [2], and then the group height was inferred from Nadler's equation - using the mean blood volume, mean sex, and mean weight.

$$
\begin{aligned}
&n_{females} = 1 \\
&n_{males} = 3 \\
&sex = \frac{n_{males}}{n_{females} + n_{males}} = \frac{3}{1+3} = 0.75 \\
&mean_{sex} = 1 \\
&mean_{weight} = 79.78 \text{ kg} \\
&mean_{bloodVolume} = 53.5540 \text{ dL} \\
&height = \left( \frac{mean_{bloodVolume} - 0.03219 \cdot mean_{weight} + 0.6041}{0.3669} \right)^{\frac{1}{3}} \\
&height = \left( \frac{53.5540 - 0.03219 \cdot 79.78 + 0.6041}{0.3669} \right)^{\frac{1}{3}} \\
&height = 1.812 \text{ m}
\end{aligned} \tag{20}
$$

#### Estimated inputs

Knowing that the density of ethanol is equal to 0.7891 g/ml or kg/L and the reported consumption of 0.51 g ethanol per kg bodyweight beer and 0.rq g per kg bodyweight beer [4], we can estimate the inputs. We assume that the 1h period of the beer was equally consumed over 4 blocks of 10 minutes to account for the blood and urine samples that was taken.

$$
\begin{aligned}
&mass_{EtOHBeer} = weight \cdot 0.51 = 79.78 \cdot 0.51 = 40.688 \text{ g} \\
&volume_{EtOHBeer} = \frac{40.688}{0.7891 \cdot 1000} = 0.0516 \text{ L} \\
&volume_{beer} = \frac{c2 \cdot v2}{c1} = \frac{0.0516 \cdot 1}{0.056} = 0.921 \text{ L} \\
&volume_{beerQuarter} = \frac{0.921}{4} = 0.230 \text{ L} \\
&mass_{EtOHWhiskey} = weight \cdot 0.51 = 79.78 \cdot 0.51 = 40.688 \text{ g} \\
&volume_{EtOHWhiskey} = \frac{40.688}{0.7891 \cdot 1000} = 0.0516 \text{ L} \\
&volume_{whiskey} = \frac{c2 \cdot v2}{c1} = \frac{0.0516 \cdot 1}{0.056} = 0.921 \text{ L}
\end{aligned} \tag{21}
$$

##### 3.3.6 Hoiseth low dose wine

#### Calculation of a mean person

The blood volume was calculated for the individuals in the group, using Nadler's equation [2], and then the group height was inferred from Nadler's equation - using the mean blood volume, mean sex, and mean weight.

$$
\begin{aligned}
&n_{females} = 1 \\
&n_{males} = 4 \\
&sex = \frac{n_{males}}{n_{females} + n_{males}} = \frac{4}{1+4} = 0.8 \\
&mean_{sex} = 1 \\
&mean_{weight} = 76.62 \text{ kg} \\
&mean_{bloodVolume} = 51.8522 \text{ dL} \\
&height = \left(\frac{mean_{bloodVolume} - 0.03219 \cdot mean_{weight} + 0.6041}{0.3669}\right)^{\frac{1}{3}} \\
&height = \left(\frac{51.8522 - 0.03219 \cdot 76.62 + 0.6041}{0.3669}\right)^{\frac{1}{3}} \\
&height = 1.793 \text{ m}
\end{aligned} \tag{22}
$$

#### Estimated inputs

Knowing that the density of ethanol is equal to 0.7891 g/ml or kg/L and the reported consumption of 0.51 g ethanol per kg bodyweight beer and 0.25 g per kg bodyweight wine [4], we can estimate the inputs. We assume that the 1h period of the beer was equally consumed over 4 blocks of 10 minutes to account for the blood and urine samples that was taken.

$$
\begin{aligned}
mass_{EtOHBeer} &= weight \cdot 0.51 = 76.62 \cdot 0.51 = 39.076 \text{ g} \\
volume_{EtOHBeer} &= \frac{39.076}{0.7891 \cdot 1000} = 0.0495 \text{ L} \\
volume_{beer} &= \frac{c2 \cdot v2}{c1} = \frac{0.0495 \cdot 1}{0.056} = 0.884 \text{ L} \\
volume_{beerQuarter} &= \frac{0.884}{4} = 0.221 \text{ L} \\
mass_{EtOHWine} &= weight \cdot 0.25 = 76.62 \cdot 0.25 = 19.155 \text{ g} \\
volume_{EtOHWine} &= \frac{19.155}{0.7891 \cdot 1000} = 0.0243 \text{ L} \\
volume_{wine} &= \frac{c2 \cdot v2}{c1} = \frac{0.0243 \cdot 1}{0.13} = 0.187 \text{ L}
\end{aligned} \tag{23}
$$

##### 3.3.7 Hoiseth medium dose wine

#### Calculation of a mean person

The blood volume was calculated for the individuals in the group, using Nadler's equation [2], and then the group height was inferred from Nadler's equation - using the mean blood volume, mean sex, and mean weight.

$$
\begin{aligned}
n_{females} &= 1 \\
n_{males} &= 4 \\
sex &= \frac{n_{males}}{n_{females} + n_{males}} = \frac{4}{1+4} = 0.8 \\
mean_{sex} &= 1 \\
mean_{weight} &= 73.54 \text{ kg} \\
mean_{bloodVolume} &= 50.9126 \text{ dL} \\
height &= \left( \frac{mean_{bloodVolume} - 0.03219 \cdot mean_{weight} + 0.6041}{0.3669} \right)^{\frac{1}{3}} \\
height &= \left( \frac{50.9126 - 0.03219 \cdot 73.54 + 0.6041}{0.3669} \right)^{\frac{1}{3}} \\
height &= 1.794 \text{ m}
\end{aligned} \tag{24}
$$

#### Estimated inputs

Knowing that the density of ethanol is equal to 0.7891 g/ml or kg/L and the reported consumption of 0.51 g ethanol per kg bodyweight beer and 0.51 g per kg bodyweight wine [4], we can estimate the inputs. We assume that the 1h period of the beer was equally consumed over 4 blocks of 10 minutes to account for the blood and urine samples that was taken.

$$
\begin{aligned}
&mass_{EtOHBeer} = weight \cdot 0.51 = 73.54 \cdot 0.51 = 37.505 \text{ g} \\
&volume_{EtOHBeer} = \frac{37.505}{0.7891 \cdot 1000} = 0.0475 \text{ L} \\
&volume_{beer} = \frac{c2 \cdot v2}{c1} = \frac{0.0475 \cdot 1}{0.056} = 0.848 \text{ L} \\
&volume_{beerQuarter} = \frac{0.848}{4} = 0.212 \text{ L} \\
&mass_{EtOHWine} = weight \cdot 0.51 = 73.54 \cdot 0.51 = 37.505 \text{ g} \\
&volume_{EtOHWine} = \frac{37.505}{0.7891 \cdot 1000} = 0.0475 \text{ L} \\
&volume_{wine} = \frac{c2 \cdot v2}{c1} = \frac{0.0475 \cdot 1}{0.13} = 0.365 \text{ L}
\end{aligned}
\tag{25}
$$

##### 3.3.8 Hoiseth low dose vodka

###### Calculation of a mean person

The blood volume was calculated for the individuals in the group, using Nadler's equation [2], and then the group height was inferred from Nadler's equation - using the mean blood volume, mean sex, and mean weight.

$$
\begin{aligned}
&n_{females} = 5 \\
&n_{males} = 0 \\
&mean_{sex} = 0 \\
&mean_{weight} = 69.84 \text{ kg} \\
&mean_{bloodVolume} = 43.0763 \text{ dL} \\
&height = \left( \frac{mean_{bloodVolume} - 0.03308 \cdot mean_{weight} + 0.1833}{0.3561} \right)^{\frac{1}{3}} \\
&height = \left( \frac{43.0763 - 0.03308 \cdot 69.84 + 0.1833}{0.3561} \right)^{\frac{1}{3}} \\
&height = 1.721 \text{ m}
\end{aligned}
\tag{26}
$$

###### Estimated inputs

Knowing that the density of ethanol is equal to 0.7891 g/ml or kg/L and the reported consumption of 0.51 g ethanol per kg bodyweight vodka and 0.25 g per kg bodyweight vodka [4], we can estimate the inputs. We assume that the 1h period of the beer was equally consumed over 4 blocks of 10 minutes to account for the blood and urine samples that was taken. We also assume that the subjects drank equal volume of non-alcoholic beverages alongside the spirits - effectively doubling the volume and halving the concentration (20 v/v%).

$$
\begin{aligned}
mass_{EtOHBeer} &= weight \cdot 0.51 = 69.84 \cdot 0.51 = 35.618 \text{ g} \\
volume_{EtOHBeer} &= \frac{35.618}{0.7891 \cdot 1000} = 0.0451 \text{ L} \\
volume_{beer} &= \frac{c2 \cdot v2}{c1} = \frac{0.0451 \cdot 1}{0.056} = 0.805 \text{ L} \\
volume_{beerQuarter} &= \frac{0.805}{4} = 0.201 \text{ L} \\
mass_{EtOHVodka} &= weight \cdot 0.25 = 69.84 \cdot 0.25 = 17.46 \text{ g} \\
volume_{EtOHVodka} &= \frac{17.46}{0.7891 \cdot 1000} = 0.0221 \text{ L} \\
volume_{vodka} &= \frac{c2 \cdot v2}{c1} = \frac{0.0221 \cdot 1}{0.20} = 0.111 \text{ L}
\end{aligned} \tag{27}
$$

##### 3.3.9 Hoiseth medium dose vodka

#### Calculation of a mean person

The blood volume was calculated for the individuals in the group, using Nadler's equation [2], and then the group height was inferred from Nadler's equation - using the mean blood volume, mean sex, and mean weight.

$$
\begin{aligned}
& n_{females} = 2 \\
& n_{males} = 3 \\
& sex = \frac{n_{males}}{n_{females} + n_{males}} = \frac{3}{2+3} = 0.6 \\
& mean_{sex} = 1 \\
& mean_{weight} = 82.18 \text{ kg} \\
& mean_{bloodVolume} = 53.1591 \text{ dL} \\
& height = \left( \frac{mean_{bloodVolume} - 0.03219 \cdot mean_{weight} + 0.6041}{0.3669} \right)^{\frac{1}{3}} \\
& height = \left( \frac{53.1591 - 0.03219 \cdot 82.18 + 0.6041}{0.3669} \right)^{\frac{1}{3}} \\
& height = 1.779 \text{ m}
\end{aligned} \tag{28}
$$

#### Estimated inputs

Knowing that the density of ethanol is equal to 0.7891 g/ml or kg/L and the reported consumption of 0.51 g ethanol per kg bodyweight beer and 0.51 g per kg bodyweight vodka [4], we can estimate the inputs. We assume that the 1h period of the beer was equally consumed over 4 blocks of 10 minutes to account for the blood and urine samples that was taken. We also assume that the subjects drank equal volume of non-alcoholic beverages alongside the spirits - effectively doubling the volume and halving the concentration (20 v/v%).

$$
\begin{aligned}
&mass_{EtOHBeer} = weight \cdot 0.51 = 82.18 \cdot 0.51 = 41.912 \text{ g} \\
&volume_{EtOHBeer} = \frac{41.912}{0.7891 \cdot 1000} = 0.0531 \text{ L} \\
&volume_{beer} = \frac{c2 \cdot v2}{c1} = \frac{0.0531 \cdot 1}{0.056} = 0.948 \text{ L} \\
&volume_{beerQuarter} = \frac{0.948}{4} = 0.237 \text{ L} \\
&mass_{EtOHVodka} = weight \cdot 0.51 = 82.18 \cdot 0.51 = 41.912 \text{ g} \\
&volume_{EtOHVodka} = \frac{41.912}{0.7891 \cdot 1000} = 0.0531 \text{ L} \\
&volume_{vodka} = \frac{c2 \cdot v2}{c1} = \frac{0.0531 \cdot 1}{0.20} = 0.266 \text{ L}
\end{aligned}
\tag{29}
$$

###### 3.3.10 Hoiseth high dose vodka

**Calculation of a mean person**

The blood volume was calculated for the individuals in the group, using Nadler's equation [2], and then the group height was inferred from Nadler's equation - using the mean blood volume, mean sex, and mean weight.

$$
\begin{aligned}
&n_{females} = 0 \\
&n_{males} = 5 \\
&mean_{sex} = 1 \\
&mean_{weight} = 80.78 \text{ kg} \\
&mean_{bloodVolume} = 54.1104 \text{ dL} \\
&height = \frac{(mean_{bloodVolume} - 0.03219 \cdot mean_{weight} + 0.6041)^{\frac{1}{3}}}{0.3669} \\
&height = \frac{(54.1104 - 0.03219 \cdot 80.78 + 0.6041)^{\frac{1}{3}}}{0.3669} \\
&height = 1.816 \text{ m}
\end{aligned}
\tag{30}
$$

**Estimated inputs**

Knowing that the density of ethanol is equal to 0.7891 g/ml or kg/L and the reported consumption of 0.51 g ethanol per kg bodyweight beer and 0.85 g per kg bodyweight whiskey [4], we can estimate the inputs. We assume that the 1h period of the beer was equally consumed over 4 blocks of 10 minutes to account for the blood and urine samples that was taken. We also assume that the subjects drank equal volume of non-alcoholic beverages alongside the spirits - effectively doubling the volume and halving the concentration (20 v/v%).

$$\begin{aligned}
mass_{EtOHBeer} &= weight \cdot 0.51 = 80.78 \cdot 0.51 = 41.120 \text{ g} \\
volume_{EtOHBeer} &= \frac{41.120}{0.7891 \cdot 1000} = 0.0521 \text{ L} \\
volume_{beer} &= \frac{c2 \cdot v2}{c1} = \frac{0.0521 \cdot 1}{0.056} = 0.930 \text{ L} \\
volume_{beerQuarter} &= \frac{0.930}{4} = 0.233 \text{ L} \\
mass_{EtOHVodka} &= weight \cdot 0.85 = 80.78 \cdot 0.85 = 68.663 \text{ g} \\
volume_{EtOHVodka} &= \frac{68.663}{0.7891 \cdot 1000} = 0.087 \text{ L} \\
volume_{vodka} &= \frac{c2 \cdot v2}{c1} = \frac{0.087 \cdot 1}{0.20} = 0.435 \text{ L}
\end{aligned} \tag{31}$$

###### 3.3.11 Wang

##### Calculation of a mean person

We first calculate the weight of the females and males using the reported mean bmi 20.9 and the mean height of 1.60 m for female and 1.72 m for males. We then calculate the blood volumes of the different participant groups using Nadler's equation [2], and the ratio of males in the participant group.

$$\begin{aligned}
weight_{females} &= height_{females} \cdot bmi = 1.60^2 \cdot 20.9 = 53.50 \text{ kg} \\
weight_{males} &= height_{males} \cdot bmi = 1.72^2 \cdot 20.9 = 61.83 \text{ kg} \\
n_{females} &= 12 \\
n_{males} &= 14 \\
sex &= \frac{n_{males}}{n_{females} + n_{males}} = \frac{14}{12+14} = 0.538 \\
vol_{bloodFemale} &= 0.3561 \cdot height^3_{female} + 0.03308 \cdot weight_{female} + 0.1833 \\
&= 0.3561 \cdot 1.60^3 + 0.03308 \cdot 53.5 + 0.1833 = 34.1 \text{ dL} \\
vol_{bloodMale} &= 0.3669 \cdot height^3_{male} + 0.03219 \cdot weight_{male} + 0.6041 \\
&= 0.3669 \cdot 1.72^3 + 0.03219 \cdot 61.83 + 0.6041 = 44.6 \text{ dL} \\
vol_{blood} &= sex \cdot vol_{bloodMale} + (1 - sex) \cdot vol_{bloodFemale} \\
&= 0.538 \cdot 44.6 + (1 - 0.538) \cdot 34.1 = 39.75 \text{ dL}
\end{aligned} \tag{32}$$

Since the ratio of males is greater than 0.5, we used a male as the average person for this study. We then calculate the height, based on the mean values, and the weight based on the mean blood volume (again using Nadler's equation) to get an approximate male representing the average person in the study.

$$\begin{aligned}
height_{male} &= \frac{height_{males} \cdot 14 + height_{females} \cdot 12}{14 + 12} = \frac{1.72 \cdot 14 + 1.60 \cdot 12}{26} = 1.665 \text{ m} \\
weight &= \frac{vol_{blood} - 0.6041 - 0.3669 \cdot height^3_{male}}{0.03219} \\
&= \frac{3.975 - 0.6041 - 0.3669 \cdot 1.665^3}{0.03219} = 52.1 \text{ kg}
\end{aligned} \tag{33}$$

##### Estimated inputs

Knowing that the density of ethanol is equal to 0.7891 g/ml or kg/L and the reported consumption of

0.72 g ethanol per kg bodyweight [5], we can estimate the inputs.

$$
\begin{aligned}
mass_{EtOH} &= weight \cdot 0.72 = 52.1 \cdot 0.72 = 37.512 \text{ g} \\
volume_{EtOH} &= \frac{37.512}{0.7891 \cdot 1000} = 0.0475377 \text{ L} \\
volume &= \frac{c2 \cdot v2}{c1} = \frac{0.0475377 \cdot 1}{0.40} = 0.1188 \text{ L}
\end{aligned} \tag{34}
$$

##### 3.3.12 Okabe 500ml Water

$$kcal = kcal_{perVol} \cdot vol = 0 \cdot 0.5 = 0 \text{ kcal} \tag{35}$$

##### 3.3.13 Okabe 500ml orange juice

$$kcal = kcal_{perVol} \cdot vol = 440 \cdot 0.5 = 220 \text{ kcal} \tag{36}$$

##### 3.3.14 Okabe 500ml milk water

$$kcal = kcal_{perVol} \cdot vol = 440 \cdot 0.5 = 220 \text{ kcal} \tag{37}$$

##### 3.3.15 Okabe 500ml orange juice and syrup

$$kcal = kcal_{perVol} \cdot vol = 660 \cdot 0.5 = 330 \text{ kcal} \tag{38}$$

##### 3.3.16 Okabe 500ml milk

$$kcal = kcal_{perVol} \cdot vol = 660 \cdot 0.5 = 330 \text{ kcal} \tag{39}$$

##### 3.3.17 Okabe 150ml Water

$$kcal_{perVol} = \frac{kcal}{vol} = \frac{0}{0.15} = 0 \text{ kcal/L} \tag{40}$$

##### 3.3.18 Okabe 150ml glucose

$$kcal_{perVol} = \frac{kcal}{vol} = \frac{67}{0.15} = 446.67 \text{ kcal/L} \tag{41}$$

##### 3.3.19 Okabe 150ml whiskey

$$kcal_{perVol} = \frac{kcal}{vol} = \frac{0}{0.15} = 0 \text{ kcal/L} \tag{42}$$

##### 3.3.20 Okabe 200ml unifrom glucose

$$kcal_{perVol} = \frac{kcal}{vol} = \frac{200}{0.2} = 1000 \text{ kcal/L} \tag{43}$$

##### 3.3.21 Okabe 400ml unifrom glucose

$$kcal_{perVol} = \frac{kcal}{vol} = \frac{200}{0.4} = 500 \text{ kcal/L} \tag{44}$$

###### 3.3.22 Okabe 600ml unifrom glucose

$$kcal_{perVol} = \frac{kcal}{vol} = \frac{200}{0.6} = 333.33 \text{ kcal/L} \tag{45}$$

###### 3.3.23 Mitchell

The drinks used in Mitchell *et al.* contained 0.5 g ethanol per kg body weight [6]. Furthermore, ethanol contains 7 kcal/g and the density of ethanol is 0.7891 g/mL. Using this information we can estimate the total volume of the ethanol in the drink.

$$\begin{aligned}
height &= \sqrt{\frac{weight}{BMI}} = \sqrt{\frac{82.66}{22.635}} = 1.771 \text{ m} \\
mass_{EtOH} &= EtOH_{dose} \cdot weight = 0.5 \cdot 82.66 = 41.33 \text{ g} \\
EtOH_{kcal} &= mass_{EtOH} \cdot EtOH_{kcalPerG} = 41.33 \cdot 7 = 289.31 \text{ kcal} \\
vol_{EtOH} &= \frac{EtOH_{kcal}}{density} = \frac{41.33}{0.7891} \cdot 0.001 = 0.052376 \text{ L}
\end{aligned} \tag{46}$$

**Beer**

For beer, it was reported that a 80 kg male was drinking a total of 409 kcal. From this, we can calculate the total volume, total kcal and total kcal/volume of the drinks, knowing that the reported mean wight was 82.66 kg:

$$\begin{aligned}
volume &= \frac{c2 \cdot v2}{c1} = \frac{0.052376 \cdot 1}{0.051} = 1.027 \text{ L} \\
total_{kcal} &= \frac{kcal}{kg} \cdot weight = \frac{409}{80} \cdot 82.66 = 422.60 \text{ kcal} \\
Beverage_{kcal} &= total_{kcal} - EtOH_{kcal} = 422.6 - 289.31 = 133.29 \text{ kcal} \\
kcal_{perVol} &= \frac{133.29}{1.027} = 129.79 \text{ kcal/L}
\end{aligned} \tag{47}$$

**Wine**

For wine, it was reported that a 80 kg male was drinking a total of 334 kcal. From this, we can calculate the total volume, total kcal and total kcal/volume of the drinks, knowing that the reported mean wight was 82.66 kg:

$$\begin{aligned}
volume &= \frac{c2 \cdot v2}{c1} = \frac{0.052376 \cdot 1}{0.125} = 0.419 \text{ L} \\
total_{kcal} &= \frac{kcal}{kg} \cdot weight = \frac{334}{80} \cdot 82.66 = 345.11 \text{ kcal} \\
Beverage_{kcal} &= total_{kcal} - EtOH_{kcal} = 345.11 - 289.31 = 55.8 \text{ kcal} \\
kcal_{perVol} &= \frac{55.8}{0.419} = 133.17 \text{ kcal/L}
\end{aligned} \tag{48}$$

**Spirit**

For spirit, it was reported that a 80 kg male was drinking a total of 297 kcal. From this, we can calculate the total volume, total kcal and total kcal/volume of the drinks, knowing that the reported mean wight was 82.66 kg:

$$
\begin{aligned}
&volume = \frac{c2 \cdot v2}{c1} = \frac{0.052376 \cdot 1}{0.051} = 0.2618 \text{ L} \\
&total_{kcal} = \frac{kcal}{kg} \cdot weight = \frac{297}{80} \cdot 82.66 = 306.88 \text{ kcal} \\
&Beverage_{kcal} = total_{kcal} - EtOH_{kcal} = 306.88 - 289.31 = 17.57 \text{ kcal} \\
&kcal_{perVol} = \frac{17.57}{0.2618} = 67.11 \text{ kcal/L}
\end{aligned} \tag{49}
$$

###### 3.3.24 Sarkola

First group of women, female1, has $n = 10$ individuals and anthropometrics, $sex = 0$, $BMI = 21.5$, estimated height 1.66 m as it was average height of women in Finland for the mean age of the participants and year of the study.

$$
weight_{female1} = BMI \cdot height^2 = 21.5 \cdot 1.66^2 = 59.24 \text{ kg} \tag{50}
$$

Second group of women, female2, has $n = 12$ individuals and anthropometrics, $sex = 0$, $BMI = 21.9$, estimated height 1.66 m as it was average height of women in Finland for the mean age of the participants and year of the study.

$$
weight_{female2} = BMI \cdot height^2 = 21.9 \cdot 1.66^2 = 60.35 \text{ kg} \tag{51}
$$

The group of men, male, has $n = 13$ individuals and anthropometrics, $sex = 1$, $BMI = 23$, estimated height 1.80 m as it was average height of men in Finland for the mean age of the participants and year of the study.

$$
weight_{male} = BMI \cdot height^2 = 23.0 \cdot 1.80^2 = 74.52 \text{ kg} \tag{52}
$$

**Calculation of a mean person**

We first calculate the blood volumes of the different participant groups using Nadler's equation [2], and the ratio of males in the participant group:

$$
\begin{aligned}
vol_{bloodFemale1} &= 0.3561 \cdot height^3_{female1} + 0.03308 \cdot weight_{female1} + 0.1833 \\
&= 37.718660055999997 \text{ dL} \\
vol_{bloodFemale2} &= 0.3561 \cdot height^3_{female2} + 0.03308 \cdot weight_{female2} + 0.1833 \\
&= 38.085848056 \text{ dL} \\
vol_{bloodMale} &= 0.3669 \cdot height^3_{male} + 0.03219 \cdot weight_{male} + 0.6041 \\
&= 51.42659600000001 \text{ dL} \\
vol_{blood} &= \frac{13 \cdot vol_{bloodMale} + 10 \cdot vol_{bloodFemale1} + 12 \cdot vol_{bloodFemale2}}{10 + 12 + 13} \\
&= \frac{13 \cdot 51.42659600000001 + 12 \cdot 38.085848056 + 10 \cdot 37.718660055999997}{35} \\
&= 42.93607214948572 \text{ dL} \\
sex &= \frac{n_{male}}{n_{total}} = \frac{13}{10 + 12 + 13} = 0.37142857142857144
\end{aligned} \tag{53}
$$

Since the ratio of males is less than 0.5, we use a female as the average person for this study. We then calculate the weight, from the mean estimated weights, and the height based on the mean blood volume (again using Nadler's equation) to get an approximate woman representing the average person in the study.

$$
\begin{aligned}
weight &= \frac{weight_{female1} \cdot 10 + weight_{female2} \cdot 12 + weight_{male} \cdot 13}{10 + 12 + 13} \\
&= \frac{59.24 \cdot 10 + 60.35 \cdot 12 + 74.52 \cdot 13}{35} = 65.30 \text{ kg} \\
height &= \left( \frac{vol_{blood} - 0.1833 - 0.03308 \cdot weight}{0.3561} \right)^{\frac{1}{3}} \\
&= \left( \frac{4.293 - 0.1833 - 0.03308 \cdot 65.3}{0.3561} \right)^{\frac{1}{3}} = 1.762 \text{ m} \\
BMI &= \frac{weight}{height^2} = \frac{65.3}{1.762^2} = 21.03 \text{ kg}/m^2
\end{aligned}
\tag{54}
$$

**Estimated inputs**

The drink used by Sarkola *et al.* contained 0.5 g ethanol per kg body weight [7]. Furthermore, the density of ethanol is 0.7891 g/mL. Using this information we can estimate the total drink volume and kcal per volume.

$$
\begin{aligned}
&mass_{EtOH} = 0.5 \cdot weight = 0.5 \cdot 65.3 = 32.65 \text{ g} \\
&vol_{EtOH} = \frac{mass_{EtOH}}{density} = 32.65 \cdot \frac{0.001}{0.7891} = 0.041376 \text{ L} \\
&volume = \frac{v1 \cdot c1}{c2} = \frac{0.041376 \cdot 1}{0.1} = 0.41376 \text{ L} \\
&Beverage_{kcal} = (volume - vol_{EtOH}) \cdot \frac{kcal}{L} = (0.41376 - 0.041376) \cdot \frac{35}{0.1} = 130.33 \text{ kcal} \\
&kcal_{perVol} = \frac{133.33}{0.41376} = 322.24 \text{ kcal/L}
\end{aligned}
\tag{55}
$$

##### 3.3.25 Jones fasting and food

The drink used by Jones *et al.* contained 0.3 g ethanol per kg body weight [8]. Furthermore, the density of ethanol is 0.7891 g/mL. Using this information we can estimate the total drink volume and kcal per volume.

$$
\begin{aligned}
&BMI = \frac{weight}{height^2} = \frac{75.8}{1.83^2} = 22.63 \text{ kg}/m^2 \\
&mass_{EtOH} = 0.3 \cdot weight = 0.3 \cdot 75.8 = 22.74 \text{ g} \\
&vol_{EtOH} = \frac{mass_{EtOH}}{density} = 22.74 \cdot \frac{0.001}{0.7891} = 0.028818 \text{ L} \\
&volume = \frac{v1 \cdot c1}{c2} = \frac{0.028818 \cdot 1}{0.2} = 0.1441 \text{ L} \\
&Beverage_{kcal} = (volume - vol_{EtOH}) \cdot \frac{kcal}{L} = (0.1441 - 0.028818) \cdot \frac{45}{0.1} = 51.877 \text{ kcal} \\
&kcal_{perVol} = \frac{51.877}{0.1441} = 360.0 \text{ kcal/L}
\end{aligned}
\tag{56}
$$

**Food**

The meal was consumed over 15 minutes and approximately 5 additional minutes before the drink was served, which gives $time_{meal} = -20$

##### 3.3.26 Javors low dose

Height for males assumed to be 1.78 m as it was average height of males in USA for the mean age of the participants and year of the study. Height for females assumed to be 1.63 m as it was average height of females in USA for the mean age of the participants and year of the study.

###### Calculation of a mean person

We first calculate the blood volumes of the different participant groups using Nadler's equation [2], and the ratio of males in the participant group.

$$\begin{aligned}
&n_{females} = 8 \\
&n_{males} = 8 \\
&sex = \frac{n_{males}}{n_{females} + n_{males}} = \frac{8}{8+8} = 0.5 \\
&vol_{bloodFemale} = 0.3561 \cdot height^3_{female} + 0.03308 \cdot weight_{female} + 0.1833 \\
&\qquad = 0.3561 \cdot 1.63^3 + 0.03308 \cdot 72.7 + 0.1833 = 41.3 \text{ dL} \\
&vol_{bloodMale} = 0.3669 \cdot height^3_{male} + 0.03219 \cdot weight_{male} + 0.6041 \\
&\qquad = 0.3669 \cdot 1.78^3 + 0.03219 \cdot 72.7 + 0.6041 = 50.1 \text{ dL} \\
&vol_{blood} = sex \cdot vol_{bloodMale} + (1 - sex) \cdot vol_{bloodFemale} \\
&\qquad = 0.5 \cdot 50.1 + (1 - 0.5) \cdot 41.3 = 45.7 \text{ dL}
\end{aligned} \tag{57}$$

Since the ratio of males is equal to 0.5, we used a male as the average person for this study. We then calculate the weight and height based on the mean blood volume (again using Nadler's equation) to get an approximate male representing the average person in the study.

$$\begin{aligned}
&height = height_{male} = 1.78 \text{ m} \\
&weight = \frac{vol_{blood} - 0.6041 - 0.3669 \cdot height^3_{male}}{0.03219} \\
&\qquad = \frac{4.57 - 0.6041 - 0.3669 \cdot 1.78^3}{0.03219} = 58.92124856166512 \text{ kg} \\
&BMI = \frac{58.92124856166512}{1.78^2} = 18.60 \text{ kg}/m^2
\end{aligned} \tag{58}$$

###### Estimated inputs

Knowing that the density of ethanol is equal to 0.7891 g/ml or kg/L and the reported consumption of 0.25 g ethanol per kg bodyweight [9], we can estimate the inputs.

$$\begin{aligned}
&mass_{EtOH} = 0.25 \cdot weight = 0.25 \cdot 72.7 = 18.175 \text{ g} \\
&conc = \frac{mass_{EtOH}}{density \cdot volume} = \frac{18.175 \cdot 0.001}{0.7891 \cdot 0.70976} = 3.245 \ \% \\
&Beverage_{kcal} = (volume - vol_{EtOH}) \cdot \frac{kcal}{L} = (0.70976 - 0.018175) \cdot \frac{35}{0.1} = 242.05 \text{ kcal} \\
&kcal_{perVol} = \frac{242.05}{0.70976} = 341.03 \text{ kcal/L}
\end{aligned} \tag{59}$$

##### 3.3.27 Javors high dose

Height for males assumed to be 1.78 m as it was average height of males in USA for the mean age of the participants and year of the study. Height for females assumed to be 1.63 m as it was average height of females in USA for the mean age of the participants and year of the study.

#### Calculation of a mean person

We first calculate the blood volumes of the different participant groups using Nadler's equation [2], and the ratio of males in the participant group.

$$\begin{aligned}
n_{females} &= 5 \\
n_{males} &= 6 \\
sex &= \frac{n_{males}}{n_{females} + n_{males}} = \frac{6}{5+6} = 0.5454 \\
vol_{bloodFemale} &= 0.3561 \cdot height^3_{female} + 0.03308 \cdot weight_{female} + 0.1833 \\
&= 0.3561 \cdot 1.63^3 + 0.03308 \cdot 71.8 + 0.1833 = 41.0 \text{ dL} \\
vol_{bloodMale} &= 0.3669 \cdot height^3_{male} + 0.03219 \cdot weight_{male} + 0.6041 \\
&= 0.3669 \cdot 1.78^3 + 0.03219 \cdot 71.8 + 0.6041 = 49.8 \text{ dL} \\
vol_{blood} &= sex \cdot vol_{bloodMale} + (1 - sex) \cdot vol_{bloodFemale} \\
&= 0.545 \cdot 49.8 + (1 - 0.545) \cdot 41.0 = 45.8 \text{ dL}
\end{aligned} \tag{60}$$

Since the ratio of males is greater than 0.5, we used a male as the average person for this study. We then calculate the weight and height based on the mean blood volume (again using Nadler's equation) to get an approximate male representing the average person in the study.

$$\begin{aligned}
height &= height_{male} = 1.78 \text{ m} \\
weight &= \frac{vol_{blood} - 0.6041 - 0.3669 \cdot height^3_{male}}{0.03219} \\
&= \frac{4.58 - 0.6041 - 0.3669 \cdot 1.78^3}{0.03219} = 59.231904044734385 \text{ kg} \\
BMI &= \frac{59.231904044734385}{1.78^2} = 18.70 \text{ kg}/m^2
\end{aligned} \tag{61}$$

#### Estimated inputs

Knowing that the density of ethanol is equal to 0.7891 g/ml or kg/L and the reported consumption of 0.5 g ethanol per kg bodyweight [9], we can estimate the inputs.

$$\begin{aligned}
mass_{EtOH} &= 0.5 \cdot weight = 0.5 \cdot 71.8 = 35.9 \text{ g} \\
conc &= \frac{mass_{EtOH}}{density \cdot volume} = \frac{35.9 \cdot 0.001}{0.7891 \cdot 0.70976} = 6.41 \ \% \\
Beverage_{kcal} &= (volume - vol_{EtOH}) \cdot \frac{kcal}{L} = (0.70976 - 0.0359) \cdot \frac{35}{0.1} = 235.85 \text{ kcal} \\
kcal_{perVol} &= \frac{235.85}{0.70976} = 322.29 \text{ kcal/L}
\end{aligned} \tag{62}$$

##### 3.3.28 Javors combined dose

This dataset is a combination of the low- and high dose data. Where, $n_{lowDose} = 16$ and $n_{highDose} = 11$. Since the data is a combination of two different data sets, we combined the simulations in the same way. In practice, this results in the following expression (Eq. (63)).

$$simCombined = \frac{16 \cdot simLow + 11 \cdot simHigh}{16 + 11} \tag{63}$$

where simCombined can be *e. g.* plasma ethanol levels.

##### 3.3.29 Frezza woman

Height for females assumed to be 1.61 m as it was average height of females in Italy for the mean age of the participants and year of the study. BMI for females assumed to be 23 $kg/m^2$. The age was assumed to be 35 years. The concentration of the wine was assumed to be 12%.
The drink used by Frezza *et al.* contained 0.3 g ethanol per kg body weight [10]. Furthermore, the density of ethanol is 0.7891 g/mL. Using this information we can estimate the total drink volume and kcal per volume.

$$\begin{aligned}
&weight = BMI \cdot height^2 = 23 \cdot 1.61^2 = 59.62 \text{ kg} \\
&mass_{EtOH} = 0.3 \cdot weight = 0.3 \cdot 59.62 = 17.886 \text{ g} \\
&vol_{EtOH} = \frac{mass_{EtOH}}{density} = \frac{17.886 \cdot 0.001}{0.7891} = 0.02267 \text{ L} \\
&volume = \frac{v1 \cdot c1}{c2} = \frac{0.02267 \cdot 1}{0.12} = 0.1889 \text{ L} \\
&Beverage_{kcal} = (0.1889 - 0.02267) \cdot \frac{5 \cdot 3.4}{0.1} = 28.25 \text{ kcal} \\
&kcal_{perVol} = \frac{28.25}{0.1889} = 149.6 \text{ kcal/L}
\end{aligned} \tag{64}$$

##### 3.3.30 Frezza men

The drink used by Frezza *et al.* contained 0.3 g ethanol per kg body weight [10]. Furthermore, the density of ethanol is 0.7891 g/mL. Height for men assumed to be 1.74 m as it was average height of men in Italy for the mean age of the participants and year of the study. BMI for men assumed to be 23 $kg/m^2$. The age was assumed to be 40 years. The concentration of the wine was assumed to be 12%. Using this information we can estimate the total drink volume and kcal per volume.

$$\begin{aligned}
&weight = BMI \cdot height^2 = 23 \cdot 1.74^2 = 69.63 \text{ kg} \\
&mass_{EtOH} = 0.3 \cdot weight = 0.3 \cdot 69.63 = 20.89 \text{ g} \\
&vol_{EtOH} = \frac{mass_{EtOH}}{density} = \frac{20.89 \cdot 0.001}{0.7891} = 0.02647 \text{ L} \\
&volume = \frac{v1 \cdot c1}{c2} = \frac{0.02647 \cdot 1}{0.12} = 0.221 \text{ L} \\
&Beverage_{kcal} = (0.221 - 0.02647) \cdot \frac{5 \cdot 3.4}{0.1} = 33.0 \text{ kcal} \\
&kcal_{perVol} = \frac{33.0}{0.221} = 149.6 \text{ kcal}
\end{aligned} \tag{65}$$

### 4 Usage of experimental data

All experimental data were digitalized from the original publications, see references in the tables below. The data was then dived into sets of estimation and validation data, the usage and division of data is highlighted in the tables below. For the estimation data, Supplementary Table 36, and the validation data, Supplementary Table 37, the following information is given, 1) the origin of the data, 2) which intervention the data corresponds to, 3) what type of data it is, and 4) how many data points (n) the data set consist of.

| Study | Intervention | Type of data | n |
|---|---|---|---|
| Kronstrand *et al.* 2019 [3] | Low dose whiskey | BAC, UAC | 25 |
| Kronstrand *et al.* 2019 [3] | Medium dose whiskey | BAC, UAC | 25 |
| Kronstrand *et al.* 2019 [3] | High dose whiskey | BAC, UAC | 25 |
| Hoiseth *et al.* 2020 [4] | Low dose beer | BAC, UAC, EtG, EtS | 53 |
| Hoiseth *et al.* 2020 [4] | Medium dose beer | BAC, UAC, EtG, EtS | 53 |
| Hoiseth *et al.* 2020 [4] | Low dose wine | BAC, UAC, EtG, EtS | 53 |
| Hoiseth *et al.* 2020 [4] | Medium dose wine | BAC, UAC, EtG, EtS | 53 |
| Hoiseth *et al.* 2020 [4] | Low dose vodka | BAC, UAC, EtG, EtS | 53 |
| Hoiseth *et al.* 2020 [4] | Medium dose vodka | BAC, UAC, EtG, EtS | 53 |
| Hoiseth *et al.* 2020 [4] | High dose vodka | BAC, UAC, EtG, EtS | 53 |
| Okabe *et al.* 2015 [11] | Water | Gastric emptying | 4 |
| Okabe *et al.* 2015 [11] | Orange juice | Gastric emptying | 7 |
| Okabe *et al.* 2015 [11] | Orange juice & syrup | Gastric emptying | 8 |
| Okabe *et al.* 2015 [11] | Milk & water | Gastric emptying | 7 |
| Okabe *et al.* 2015 [11] | Milk | Gastric emptying | 8 |
| Okabe *et al.* 2017 [12] | Glucose solution 200ml | Gastric emptying | 8 |
| Okabe *et al.* 2017 [12] | Uniform glucose 400ml | Gastric emptying | 10 |
| Okabe *et al.* 2017 [12] | Glucose solution 600ml | Gastric emptying | 11 |
| Okabe *et al.* 2022 [13] | Water | Gastric emptying | 10 |
| Okabe *et al.* 2022 [13] | Glucose solution | Gastric emptying | 10 |
| Okabe *et al.* 2022 [13] | Whiskey | Gastric emptying | 10 |
| Mitchell *et al.* 2014 [6] | Beer | BAC | 11 |
| Mitchell *et al.* 2014 [6] | Wine | BAC | 12 |
| Mitchell *et al.* 2014 [6] | Spirit blend | BAC | 12 |
| Jones *et al.* 1997 [14] | Spirit blend | BAC | 13 |
| Jones *et al.* 1997 [14] | Spirit blend & food | BAC | 13 |
| Javors *et al.* 2016 [9] | Spirit blend low dose | BrAC | 7 |
| Javors *et al.* 2016 [9] | Spirit blend low dose | PEth | 7 |
| Javors *et al.* 2016 [9] | Spirit blend high dose | BrAC | 7 |
| Javors *et al.* 2016 [9] | Spirit blend high dose | PEth | 7 |
| Javors *et al.* 2016 [9] | Spirit blend combined doses | PEth | 13 |
| Sarkola *et al.* 2002 [7] | Spirit blend | Acetate | 4 |
| Sarkola *et al.* 2002 [7] | Spirit blend | BAC | 4 |
| Frezza *et al.* 1990 [10] | Wine Female | BAC | 13 |
| Frezza *et al.* 1990 [10] | Wine Male | BAC | 12 |

Supplementary Table 36: **Estimation data**. Detailing the origin of the data, which intervention the data corresponds to, what type of data it is, and how many data points (n) the data set consist of.

| **Study** | **Intervention** | **Type of data** | **n** |
|---|---|---|---|
| Wang *et al.* 2022 [5] | Spirit | BAC | 7 |
| Wang *et al.* 2022 [5] | Spirit | EtG | 7 |
| Wang *et al.* 2022 [5] | Spirit | EtS | 7 |

Supplementary Table 37: **Validation data**. Detailing the origin of the data, which intervention the data corresponds to, what type of data it is, and how many data points (n) the data set consist of.

### 5 Changelog of rejected model structures

As the modelling cycle is an iterative process multiple model versions has been rejected along the project. Following these versions are listed, with a short explanation of what they tried to improve. These model files are available in the provided code for this publication.

**Version 1** - This version, named alcohol_model_UAC_EtG_EtS_1.txt in the previous model files, introduces the creation and elimination of EtG, EtS, and UAC into the model structure. It also introduced the "kcal_remain" state to keep track of the remaining kcal from the last drink and make the gastric emptying rate better.

**Version 2** - This version, named alcohol_model_UAC_EtG_EtS_2.txt in the previous model files, removes a logic gate from EtG and Ets and adds a scaling factor.

**Version 3** - This version, named alcohol_model_UAC_EtG_EtS_3.txt in the previous model files, tests EtG and EtS to behave according to mass action dynamics as Michaelis Menten kinetics were introduce directly in version 1. The faulty unit conversion factor (1000) is also removed.

**Version 4** - This version, named alcohol_model_UAC_EtG_EtS_4.txt in the previous model files, reintroduced Michaelis Menten kinetics for the synthesis of EtG and Ets. Furthermore, greater than (gt) check is introduce to avoid the noisy simulations were the concentration ends up negative.

**Version 5** - This version, named alcohol_model_UAC_EtG_EtS_5.txt in the previous model files, identifies a potential limitation with the alcoholic calories not being considered and therefore introduced the state "KcalEtOH_Liquid" and a relating remain state. It also tries to scale the rate of gastric emptying linearly and removes the Michaelis Menten dynamics.

**Version 6** - This version, named alcohol_model_UAC_EtG_EtS_6.txt in the previous model files, reintroduced Michaelis Menten dynamics on the elimination of EtG and EtS.

**Version 7** - This version, named alcohol_model_UAC_EtG_EtS_7.txt in the previous model files, reintroduces the Michaelis Menten dynamics for the gastric emptying, makes the elimination of EtG and EtS act according to mass action dynamics again, and removes EtOH kcal in the gastric emptying dynamic. It also introduces the liver as a separate compartment to allow for delay dynamics for the EtOH elimination and synthesis of EtG and EtS.

**Version 8** - This version, named alcohol_model_UAC_EtG_EtS_8.txt in the previous model files, reintroduce the EtOH calories to be part of the gastric emptying dynamics.

**Version 9** - This version, named alcohol_model_UAC_EtG_EtS_9.txt in the previous model files, introduces *total body water* (TBW) as a compartment to distribute the ethanol into. In compliance with this, concentration gradients were added to the reactions between different concentration compartments. The naming logic was updated to use "understandable" names for parameters and reaction. The stomach states was changed to use litre (L) instead of deciliter (dL).

**Version 10** - This version, named alcohol_model_UAC_EtG_EtS_10.txt in the previous model files, tried out a new "memory" of previously consumed calories. It also introduces a maximum slowing effect of the aggregated effect of kcal and $kcal_{EtOH}$ on the gastric emptying rate.

**Version 11** - This version, named alcohol_model_UAC_EtG_EtS_11.txt in the previous model files, splits the total volume of blood into a venous and arterial compartment.

**Version 12** - This version, named alcohol_model_UAC_EtG_EtS_12.txt in the previous model files, removed the $mem_{kcal}$ states as the gastric emptying event was faulty and the interactions did work properly. Arteriole and venous blood was renamed to Central and Peripheral to better represent their intended role.

**Version 13** - This version, named alcohol_model_UAC_EtG_EtS_13.txt in the previous model files, completes the Central-Peripheral renaming change, it removes the unused kcal_emptying reaction, it removes the intestines elimination reaction as it had no overall effect, it introduce personalization of the liver volume, and it introduces an intermediate state to try to slow down the UAC dynamics.

**Version 14** - This version, named alcohol_model_UAC_EtG_EtS_14.txt in the previous model files, introduces an additional homologue of the ADH enzyme.

**Version 15** - This version, named alcohol_model_UAC_EtG_EtS_15.txt in the previous model files, reworks the urine dynamics to assume a constant bladder volume and instead empties the concentration with an urination event. The model now describes the UAC with diffusion and has no elimination reaction from the bladder as the event instead removes the ethanol. The second homologue of ADH is removed as it was not effective.

**Version 16** - This version, named alcohol_model_UAC_EtG_EtS_16.txt in the previous model files, reworks the urine dynamics to consider the ethanol mass and liquid volume separately. Tissue EtG and EtS are removed and introduced to the urine compartment.

**Version 17** - This version, named alcohol_model_UAC_EtG_EtS_17.txt in the previous model files, introduce uptake of drunken volumes into a $Tissue_{extra-water}$ state that feeds the urine production, drinking more leads to more urine produced. Furthermore, the urine ethanol exchange is now compared to the plasma concentration instead of the blood concentration scaled with a fraction - the fraction represented the water content in blood, *i. e.* plasma, and is instead moved to the $V_{Plasma-Peripheral}$ variable.

**Version 18** - This version, named alcohol_model_UAC_EtG_EtS_18.txt in the previous model files, introduces the effect of vasopressin - regulating the urine produced and the permeability of the kidneys. In practice, producing more urine when ethanol is present, which inhibits vasopressin production, and reducing the exchange from the kidneys to plasma as ethanol increases.

**Version 19** - This version, named alcohol_model_UAC_EtG_EtS_19.txt in the previous model files, introduces the possibility for urine EtG and EtS to return from the kidneys to plasma dependent on the concentration difference and the presence of vasopressin.

**Version 20** - This version, named alcohol_model_UAC_EtG_EtS_20.txt in the previous model files, tries out a new version of kcal delay effect in the rate of gastric emptying, a sigmoid curve, in an effort to get EtOH kcal to be more impactful.

**Version 21** - This version, named alcohol_model_UAC_EtG_EtS_21.txt in the previous model files, tries an alternative kcal delay effect in the rate of gastric emptying, a hill equation.

**Version 22** - This version, named alcohol_model_UAC_EtG_EtS_22.txt in the previous model files, adds the effect of gastric ADH.

**Version 23** - This version, named alcohol_model_UAC_EtG_EtS_23.txt in the previous model files, removes the clearance of EtG and EtS from the blood.

**Version 24** - This version, named alcohol_model_UAC_EtG_EtS_24.txt in the previous model files, tries to describe the ADH dynamics as a mass action since the model identifiability analysis showed that Km could be removed - making the Michaelis Menten reaction a constant. The model could fit data with a mass action reaction. However, the fit was worse. This version was dropped since the biological interpretation is more logical with the Michaelis Menten reaction and a constant rate has an explanation in the enzyme is being fully saturated all the time.

**Version 25** - This version, named alcohol_model_UAC_EtG_EtS_25.txt in the previous model files, tries to remove the outflow from the food compartment in the stomach (since the parameter could be a fairy small value). The decrees in model performance was deemed to great to justify this removal.

**Version 26** - This version, named alcohol_model_UAC_EtG_EtS_26.txt in the previous model files, tries to make the gastric emptying dynamic a hill equation to have greater variation in the emptying rate.

**Version 27** - This version, named alcohol_model_UAC_EtG_EtS_27.txt in the previous model files, introduces a "greater than" expression on the synthesis of EtG and EtS to prevent the production of EtG and EtS at very low BAC levels.

**Version 28** - Final version, changes the "greater than" expression of the EtG and EtS synthesis to a delay state to improve the model stability.
